# Opposing effects of systemic and pancreas-specific inhibition of BCKDK on pancreatic carcinogenesis

**DOI:** 10.1101/2025.06.10.658925

**Authors:** Michael C. Noji, Christina Demetriadou, Madelyn Landis, Jennifer Pennise, Laura V. Pinheiro, Alison Jaccard, Adam Chatoff, Jack Drummond, Kevin Guo, Romie Azor, Maggie R. Robertson, Ryo Kawakami, Mariola M. Marcinkiewicz, John W. Tobias, Nathaniel W. Snyder, David Feldser, Zoltan Arany, Kathryn E. Wellen

## Abstract

Branched-chain amino acid (BCAA) metabolism is perturbed in patients with pancreatic cancer, but the contribution of systemic or pancreas-intrinsic BCAA catabolism to pancreatic carcinogenesis is unclear. We show here that pancreas-specific loss of DBT, the E2 subunit of the branched-chain keto-acid dehydrogenase (BCKDH) complex required for BCAA oxidation, strikingly exacerbates premalignant pancreatic intraepithelial neoplasia (PanIN) lesions in KC (*p48-Cre*;*Kras^LSL-G12D/+^*) mice. However, deletion of upstream enzyme BCAT2 neither phenocopied nor rescued loss of DBT in KC mice, ruling out involvement of both upstream and downstream metabolites as mediators of PanIN promotion. Instead, we observed that DBT deficiency led to loss of the kinase BCKDK, a negative regulator of the BCKDH complex, and that, remarkably, pancreas-specific loss of BCKDK phenocopied DBT deficiency in accelerating PanIN formation. These data thus support a model in which pancreas BCKDK restrains tumorigenesis. In contrast, systemic treatment of KC mice with the BCKDK inhibitor BT2, which inhibits BCKDH phosphorylation across many tissues except the pancreas, reduced PanIN formation and preserved normal acinar area. Together the data reveal the promotion of BCAA catabolism systemically, but not within the pancreas, as a promising intervention strategy to suppress tumor initiation.

## Introduction

Pancreatic cancer is the 4^th^ leading cause of cancer deaths worldwide and predicted to become the second leading cause by 2030^1^. Pancreatic ductal adenocarcinoma (PDA), the most common type of pancreatic cancer, has a median 5-year survival rate of 13%^2^. The incidence of pancreatic cancer has continuously risen in recent decades^3^. Although surgical resection of early-stage PDA can be curative, 80-85% of patients are diagnosed at more advanced stages^1^, for which resection is not an option and standard-of-care chemotherapy and radiotherapy produce only modest increases in survival. As such, delineating mechanisms to prevent PDA carcinogenesis or intercept it at early stages is of high importance. Previous studies have linked pancreatic cancer development with metabolic diseases. For instance, factors such as obesity^4,5^, diabetes^6,7^, and chronic pancreatitis^8–10^ confer elevated risk for developing pancreatic cancer. Mechanistically, elevated insulin levels and insulin receptor signaling in acinar cells^11–13^ and local production of cholecystokinin^14^ have been found to contribute to obesity-linked pancreatic tumorigenesis in mouse models. Moreover, some evidence suggests that nutrient intake and prandial hormone responses may contribute to PDA risk^15,16^, but the mechanistic roles of specific nutrients to carcinogenesis remains poorly understood.

Branched-chain amino acid (BCAA) metabolism has garnered substantial interest for its potential roles in pancreatic tumorigenesis. The BCAAs (isoleucine, leucine, and valine) are essential amino acids that all mammals must receive from their diet and are abundant in meat, dairy, and legumes^17^. Once in cells, BCAAs are used for protein synthesis or can be transaminated by the branched-chain amino acid aminotransferase (BCAT) enzymes to their respective branched-chain keto acids (BCKAs), which can then enter the rate limiting step of the BCAA catabolic pathway, the branched-chain keto-acid dehydrogenase (BCKDH) complex (Fig. 1a). The BCKDH complex consists of several components, including DBT, the E2 central subunit of the complex. Elevated plasma levels of BCAAs have been observed in individuals who later develop PDA^18^, even 10 years prior to diagnosis^19^ and levels of BCAAs have been found to be higher in PDA tumors than adjacent healthy tissue^20,21^. Moreover, a positive association between BCAA intake and PDA development risk was also recently reported^22^. Concordantly, in mice, a low BCAA diet reduced plasma BCAA levels and was protective against PanIN formation^23^ while a high BCAA diet increased plasma BCAA levels and promoted PanIN progression^24^, suggesting that BCAA availability might influence PDA initiation. The mitochondrial BCAT2 enzyme has been found to be protective in murine models of PDA^23,25^, but whether this is explicitly through mitochondrial BCKA catabolism is not fully understood.

**Figure 1.**
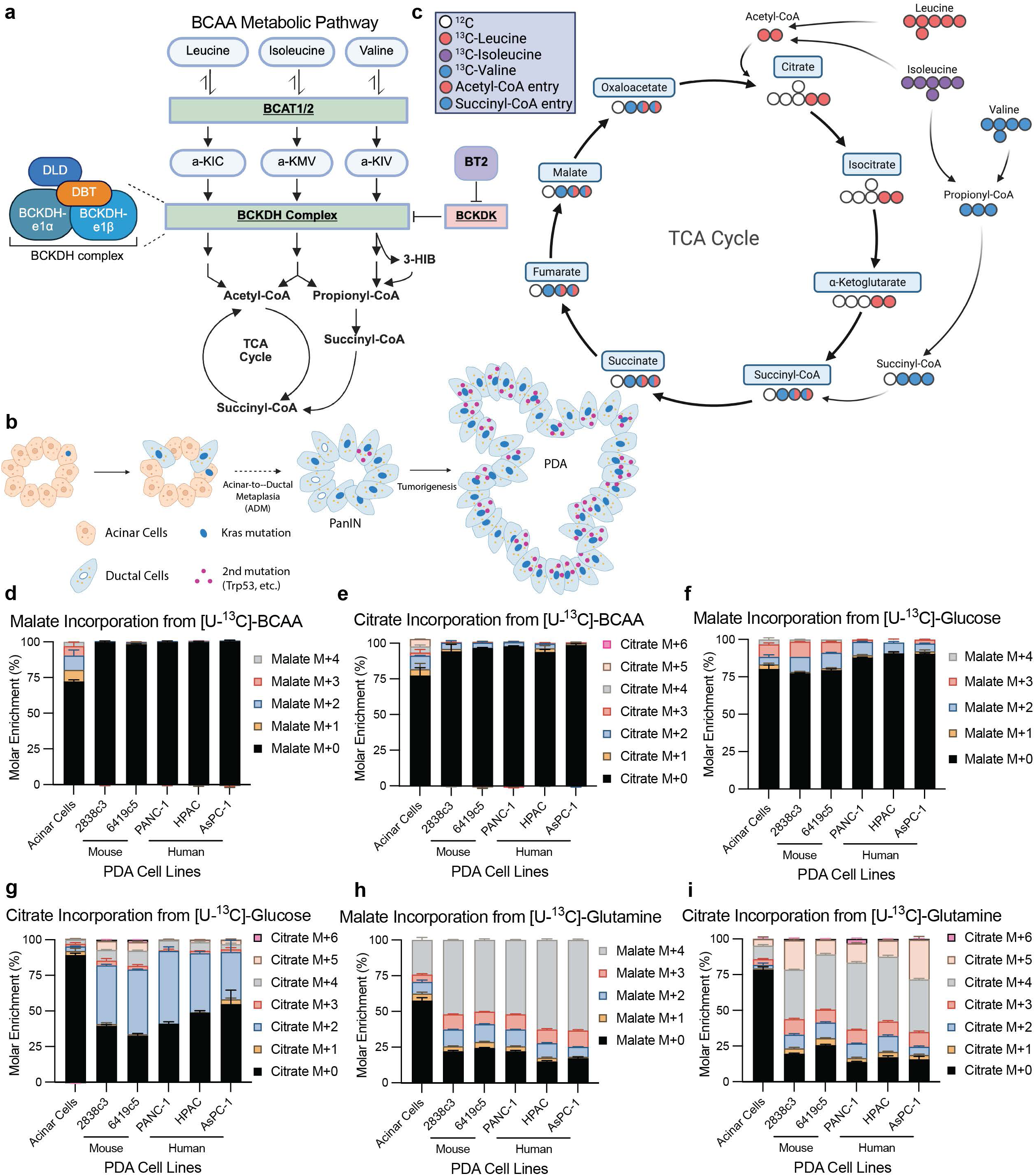
BCAAs feed the TCA cycle in healthy pancreatic acinar cells but not in PDA cells. **a**, Pathway of BCAA catabolism, highlighting key enzymes (BCAT2, BCKDH Complex including DBT, BCKDK) and a pharmacological BCKDK inhibitor, BT2. **b**, Diagram depicting PanIN formation via *KRAS* mutation following acinar-to-ductal metaplasia (ADM), followed by a second genetic mutation like *Trp53* that leads to the development of frank PDA. **c**, Tracing schematic for [U-^13^C]-BCAA incorporation into TCA cycle intermediates showing entry of labelled carbons can via acetyl-CoA (red circles) and succinyl-CoA (blue circles). **d-i**. Molar percent enrichment (MPE) of malate and citrate in a panel of PDA murine and human PDA cell lines and primary mouse acinar cells after 6-hour incubation of either [U-^13^C]-BCAAs (valine, isoleucine, leucine), [U-^13^C]-glutamine, or [U-^13^C]-glucose, measured by GC-MS (n=3 biological replicates per cell line). MPE refers to the % of each isotopologue relative to the total metabolite pool after correcting for ^13^C natural abundance. Data are presented as mean ± s.e.m.

A remarkable feature of the normal pancreas is its high use of BCAAs, both to support the substantial protein synthesis that is a function of the exocrine pancreas and for oxidation. We have demonstrated through steady-state *in vivo* infusion studies that BCAAs are a major substrate contributing to the TCA cycle in mouse pancreata^26^, contrasting their relatively minor contribution in other tissues. Consistently, BCAAs contribute prominently to acetyl-CoA production in pancreatic acinar cells, and loss of the acetyl-CoA producer ATP-citrate lyase impairs PanIN formation^27^. Since BCAAs are a major supplier to acetyl-CoA pools and the TCA cycle in the normal pancreas, we hypothesized that BCAA catabolism may also directly contribute to PanIN formation.

Here, using genetic loss-of-function approaches, we find in a murine model of PDA initiation (*p48-Cre; Kras^LSL-G12D/+^;* herein referred to as KC)^28^ that contrary to expectations, loss of the DBT in the pancreas exacerbates PanIN formation. This is linked to an unexpected suppression of BCKDK protein level and activity, and we find that pancreas-specific loss of BCKDK also promotes PanIN formation in the KC model. Pancreas-intrinsic BCKDK demonstrates a PanIN-restraining role for pancreas-intrinsic BCKDK, likely independent of its canonical role in regulation of BCKDH. Conversely, systemic inhibition of BCKDK lowers circulating availability of BCAAs and BCKAs and suppresses PanIN formation. The data reveal strikingly opposing effects of systemic and pancreas-specific inhibition of BCKDK on pancreatic carcinogenesis and suggest a novel strategy to reduce tumor formation in individuals at elevated risk.

## Results

### BCAAs feed the TCA cycle in healthy pancreatic acinar cells but not in PDA cells

Pancreatic ductal adenocarcinoma (PDA) can arise from pancreatic acinar cells that undergo acinar-to-ductal metaplasia or ADM (Fig. 1b). We first sought to quantify BCAA oxidation in PDA cells versus primary pancreatic acinar cells. For this, we evaluated the contribution of [U-^13^C]-glucose, [U-^13^C]-glutamine, and [U-^13^C]-BCAAs (leucine, isoleucine, and valine) to TCA cycle intermediates across a panel of human and murine PDA cell lines and primary acinar cells freshly isolated from murine pancreata (Fig. 1c and Extended Data Fig. 1a,b). Concordant with prior work^26^, labeling of BCAAs into malate and citrate was clear in non-neoplastic acinar cells (Fig. 1d,e). However, in PDA cells, minimal labeling from BCAAs was detected, suggesting little use of BCAAs to feed the TCA cycle in established PDA cells. In contrast, extensive labeling from glucose and glutamine was observed in all cells (Fig. 1f-i). Pool sizes for malate and citrate (Extended Data Fig. 1c-h) were similar across the cell lines. These data indicate that the contribution of BCAA catabolism to the TCA cycle is minimal in established PDA cells, but that BCAAs play a major role in contributing carbon to the TCA cycle in healthy pancreatic acinar cells, consistent with prior studies^26,27,29^. These data encouraged us to test if this loss of BCAA catabolism in PDA cells contributes to tumor formation.

### Pancreas DBT knockout is well tolerated and results in local elevation of BCAAs and BCKAs

We first aimed to determine if BCAA oxidation is essential for normal pancreas function. *Dbt*^fl/fl^ animals^26^ were crossed to *Ptf1a* (also known as *P48*)-Cre to generate DBT pancreas specific knockout mice (DBT-pKO) (Fig. 2a and Extended Data Fig. 2a-c). Pancreas labeling of acetyl-CoA and succinyl-CoA from gavaged [U-^13^C]-BCAAs was ablated in the absence of DBT (Fig. 2b,c and Extended Data Fig. 2d-h), confirming functional ablation of BCAA catabolism. Propionyl-CoA M+3 labeling was suppressed by about 60% (Fig. 2d); residual labeling may reflect potential uptake of the valine catabolism intermediate, 3-HIB, secreted by other cell types with intact BCAA catabolism. 3-HIB can be detected in circulation within minutes of providing [U-^13^C]-BCAAs to animals^26^. Fasted plasma BCAA and BCKA levels were unchanged in DBT-pKO animals (Fig. 2e), but a 2-3-fold increase in levels of both BCAAs and BCKAs was observed in the pancreas (Fig. 2f). Abundance of TCA cycle intermediates was well maintained in DBT-pKO animals (Fig. 2g), suggesting that even though BCAAs are major contributors to the TCA cycle in the healthy pancreas, there is likely metabolic compensation in the absence of BCAA catabolism. Thus, DBT deficiency blocks BCAA catabolism in the pancreas and leads to elevated tissue levels of BCAAs and BCKAs.

**Figure 2.**
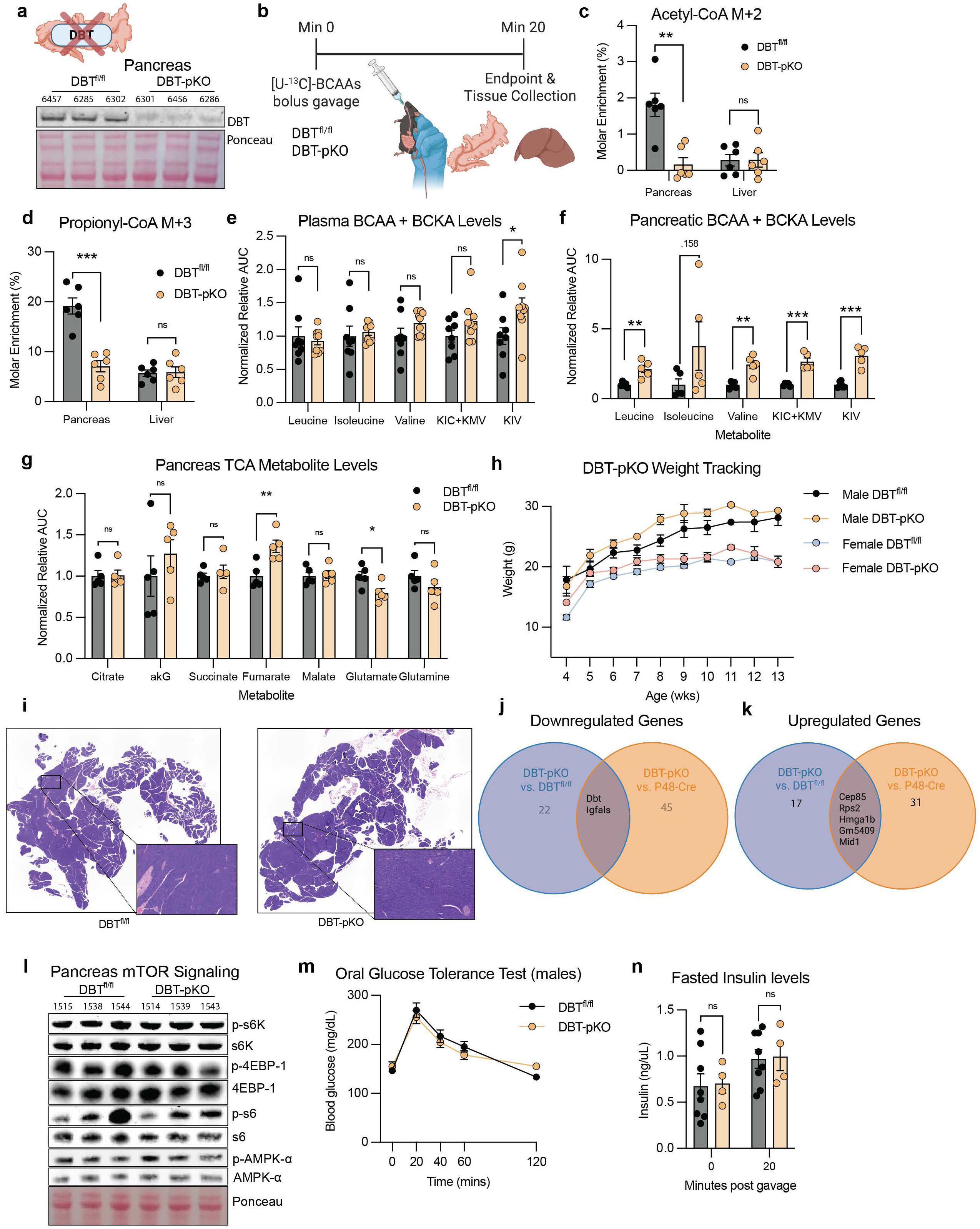
Pancreas DBT knockout is well tolerated and results in local elevation of BCAAs and BCKAs. **a,** Western blot for DBT protein in pancreata of 8-10-week-old DBT^fl/fl^ and DBT-pKO mice (n=3 DBT^fl/fl^ and n=3 DBT-pKO; Ponceau stain is the loading control). **b**, Schematic diagram of bolus [U-^13^C]-BCAA oral gavage. **c,d,** Acetyl-CoA M+2 and propionyl-CoA M+3 labeling measured in animal pancreata and liver 20 mins after bolus gavage of [U-^13^C]-BCAAs (n=6 DBT^fl/fl^ and n=6 DBT-pKO). **e**, LC-MS plasma relative measurements of BCAAs and BCKAs in 5-hour fasted animals (n=8 DBT^fl/fl^ and n=9 DBT-pKO). **f,** LC-MS pancreata relative measurements of BCAAs and BCKAs in fed animals (n=5 DBT^fl/fl^ and n=5 DBT-pKO). **g**, LC-MS relative measurements of pancreatic TCA metabolites in fed mice (n=5 DBT^fl/fl^ and n=5 DBT-pKO). **h**, Mouse weight recorded weekly in the afternoon (1:00-3:00 PM) throughout the experiment (n=12 male DBT^fl/fl^, n=11 male DBT-pKO, n=16 female DBT^fl/fl^, and n=14 female DBT-pKO). **i**, Representative H+E-stained images of pancreata from 10-week-old mice. **j,k**, Commonly downregulated and upregulated genes from bulk RNA-seq comparing DBT-pKO vs. DBT^fl/fl^ and DBT-pKO vs. P48-Cre (n=2 per experimental group). Cutoffs for significance: (abs(log2ratio)) > 1.5. and (FDR) adjusted p-value of ≤ 0.05. **l**, Western blot for mTORC1 signaling proteins in mouse pancreata (n=3 DBT^fl/fl^ and n=3 DBT-pKO; Ponceau stain is the loading control). **m**, Orally administered glucose tolerance tests (OG-GTTs) in 10-week-old male animals after a 5-hour morning fast from 7 AM-12 PM (n=9 DBT^fl/fl^ and n=14 DBT-pKO). **n**, Insulin levels measured at 0 and 20 mins post oral gavage from animals in panel m (n=8 DBT^fl/fl^ and n=4 DBT-pKO). Data are presented as mean ± s.e.m. Representative images depict animals closest to the mean of each group. For comparing two groups, a two-tailed Student’s t-test was used for each individual tissue or metabolite plotted, with significance defined as *p<0.05, **p<0.01 and ***p<0.001. **c**, pancreas acetyl-CoA M+2 *P*=0.0012. **d**, pancreas propionyl-CoA M+2 *P*=0.0001. **e**, KIV *P*=0.0373. **f**, Leucine *P*=0.0033; Isoleucine *P*=0.1581; Valine *P*=0.0016; KIC + KMV *P*=0.0003; KIV *P*=0.0002. **g**, Fumarate *P*=0.0068; Glutamate *P*=0.0272.

We next examined the phenotype of DBT-pKO animals. Total body weight (Fig. 2h), forelimb size (Extended Data Fig. 2i), and pancreas histology (Fig. 2i) were comparable between genotypes. RNA-sequencing analyses comparing DBT-pKO pancreata to either DBT^fl/fl^ littermate controls or P48-Cre;Dbt^+/+^ controls revealed minimal transcriptional changes in the absence of DBT (Fig. 2j,k and Extended Data Fig. 2j). Additionally, targeted qPCR analysis showed no substantial changes in expression of genes involved in normal exocrine or endocrine pancreatic function (Extended Data Fig. 2k,l). Since leucine is a known activator of signaling through the mammalian target of rapamycin complex 1 (mTORC1)^30^, and given that we observed increased levels of leucine in the DBT-pKO pancreas tissue (Fig. 2f), we next sought to examine signaling in DBT-pKO pancreata. We observed no differences in mTORC1 signaling, including p-S6k, p-4EBP-1, p-S6, or p-AMPK-α (Fig. 2l). Neither males (Fig. 2m,n) nor females (Extended Data Fig. 2m,n) showed differences between genotypes in glucose tolerance or insulin levels after receiving a glucose bolus. Thus, the data indicate that in unstressed mice, loss of BCAA catabolism within the pancreas results in minimal phenotypic changes.

### PanIN formation is accelerated in DBT-pKO mice

Next, to assess the effects of genetic loss of pancreatic BCAA catabolism on tumor initiation, *DBT^ff/ff^* animals were crossed to KC (*p48-Cre*;*Kras^LSL-G12D/+^)* animals to generate *KC;DBT^fl^*^/fl^ (KC;DBT-pKO) mice. The KC mouse model develops premalignant PanIN lesions, with slow progression to PDA, with survival typically over 12 months^31^. KC;DBT-pKO body weight did not significantly differ from control animals (Extended Data Fig. 3a). To synchronously induce acinar-to-ductal metaplasia (ADM) and PanIN formation in these mice, we leveraged a cerulein-dependent pancreatitis model^32^. Cerulein is a cholecystokinin analogue that promotes inflammation and pancreatitis, triggering ADM that progresses to PanIN in the presence of mutant KRAS (Fig 3a). Unexpectedly, 8-week-old cerulein-treated KC;DBT-pKO animals had significantly heavier and more rigid pancreata than WT controls at 9 days post-cerulein injection (Extended Data Fig. 3b), as well as more neoplastic area (Fig. 3b), with a corresponding loss of normal acinar tissue (Fig. 3c). Histopathological scoring revealed that the DBT deficient animals had both a greater number of and more advanced PanINs compared to KC controls (Fig. 3d). Moreover, both ADM and inflammation features were elevated in the KC;DBT-pKO animals (Fig. 3e,f), based on multiparametric analysis (Extended Data Fig. 3c,d). To test if this phenotype requires pancreatitis, we also examined KC;DBT-pKO mice in the absence of cerulein at 8 and 12 weeks of age, similarly finding greater loss of normal acinar area and increased presence of PanIN lesions (Fig. 3g-i and Extended Data Fig. 3e-k). Altogether, these data show that pancreatic DBT deficiency exacerbates KRAS^G12D^-driven PanIN formation.

**Figure 3.**
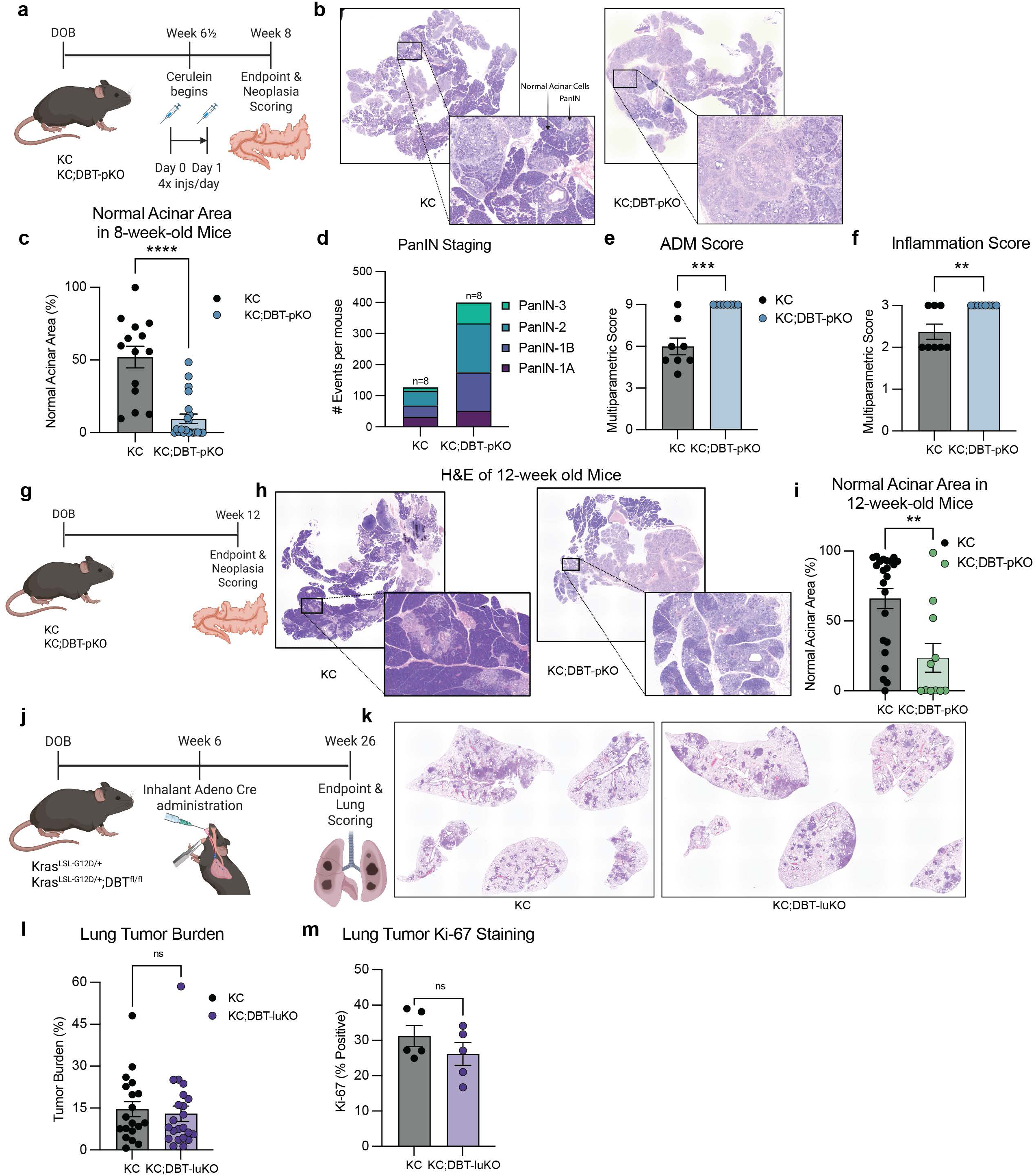
DBT loss accelerates KRAS^G12D^-driven tumor initiation in the pancreas, but not in the lungs. **a,** Schematic and timeline for tumor initiation assays for P48-Cre; Kras^LSL-G12D/+^; (KC) mice with induction of pancreatitis via intraperitoneal cerulein administration. **b**, Representative HCE images of KC and KC;DBT-pKO pancreata after induction of pancreatitis at 6 ½ weeks and collected at an 8-week timepoint. Normal acinar cells and PanIN are indicated with black arrows. **c**, Normal acinar area was calculated by the total area of healthy acinar tissue relative to total pancreas area for each animal (n=14 KC and n=21 KC;DBT-pKO). Values are measured using FIJI. **d**, Histopathological scoring of the number and severity of PanIN lesions (n=8 KC and n=8 KC;DBT-pKO. 8 animals from **c** closest to the mean within each group were chosen). **e,f**, Histopathological scoring of Acinar-to-Ductal Metaplasia (ADM) and Inflammation as graded on a multiparametric scale. Scoring is broken down in Extended Data Fig. 4c,d. **g**, Schematic and timeline for tumor initiation assays for KC mice collected at 12 weeks of age. **h**, Representative HCE images of KC and KC;DBT-pKO pancreata. **i**, Normal acinar area relative to total pancreas area of animals harvested at 12 weeks old (n=23 KC and n=11 KC;DBT-pKO). **j**, Timeline for tumor initiation assays for inhalant Ad:CMV-Cre Kras^LSL-G12D/+^ (KC) initiation model of lung adenocarcinoma. **k**, Representative HCE images of lung lobes 20 weeks after inhalant Cre delivery at endpoint. **l**, Lung tumor burden is calculated by normalizing the area of tumor burden to total lung area (n=19 KC and n=22 KC;DBT-luKO). **m**, Quantification of Ki-67 positivity in tumor cells is calculated as a % (n=5 KC and n=5 KC;DBT-luKO. 5 animals from **l** closest to the mean within each group were chosen). Data are presented as mean ± s.e.m. Representative images depict animals closest to the mean of each group. For comparing two groups, a two-tailed Student’s t-test was used, with significance defined as **p<0.01, ***p<0.001 and ****p<0.0001. **c**, Normal acinar area *P*<0.0001. **e**, ADM score *P*=0.0002. **f**, Inflammation score *P*=0.0042. **i**, Normal acinar area *P*=0.0019.

### Acceleration of tumor formation by DBT deletion is not seen in lung adenocarcinoma

KRAS mutations are also common in lung adenocarcinoma, present in up to 30% of all lung tumors^33,34^. To understand if loss of DBT exerts common effects to promote carcinogenesis in a KRAS-driven lung tumor model, we injected *Kras^LSL-G12D/+^;Dbt^ff/ff^* and *Kras^LSL-G12D/+^* control animals with inhalant Ad:CMV-Cre to promote spontaneous lung adenocarcinoma in the presence and absence of DBT (DBT-luKO) (Fig. 3j). Lung tumors formed in all animals, confirming Cre-delivery (Fig. 3k and Extended Data Fig. 3l). However, there was no difference in tumor burden with loss of DBT in the lung (Fig. 3l), in sharp contrast to our observations in pancreas. Proliferation rates measured by Ki-67 staining also show no difference in DBT-luKO animals (Fig. 3m and Extended Data Fig. 3m). Thus, loss of DBT has distinct effects in lung and pancreatic tumorigenesis, with minimal effects observed in lung.

### Loss of pancreatic BCAA catabolism is not sufficient to exacerbate PanIN formation

We next investigated the mechanism through which DBT deficiency promotes PanIN formation, hypothesizing that the loss of BCAA oxidation within the pancreas contributes to tumor formation. To test this, we generated pancreas-specific BCAT2 knockout animals (BCAT2-pKO) (Fig. 4a and Extended Data Fig. 4a), which, like DBT-pKO animals, have reduced pancreatic BCAA oxidation, and which we therefore hypothesized, would phenocopy the DBT-pKO animals.

**Figure 4.**
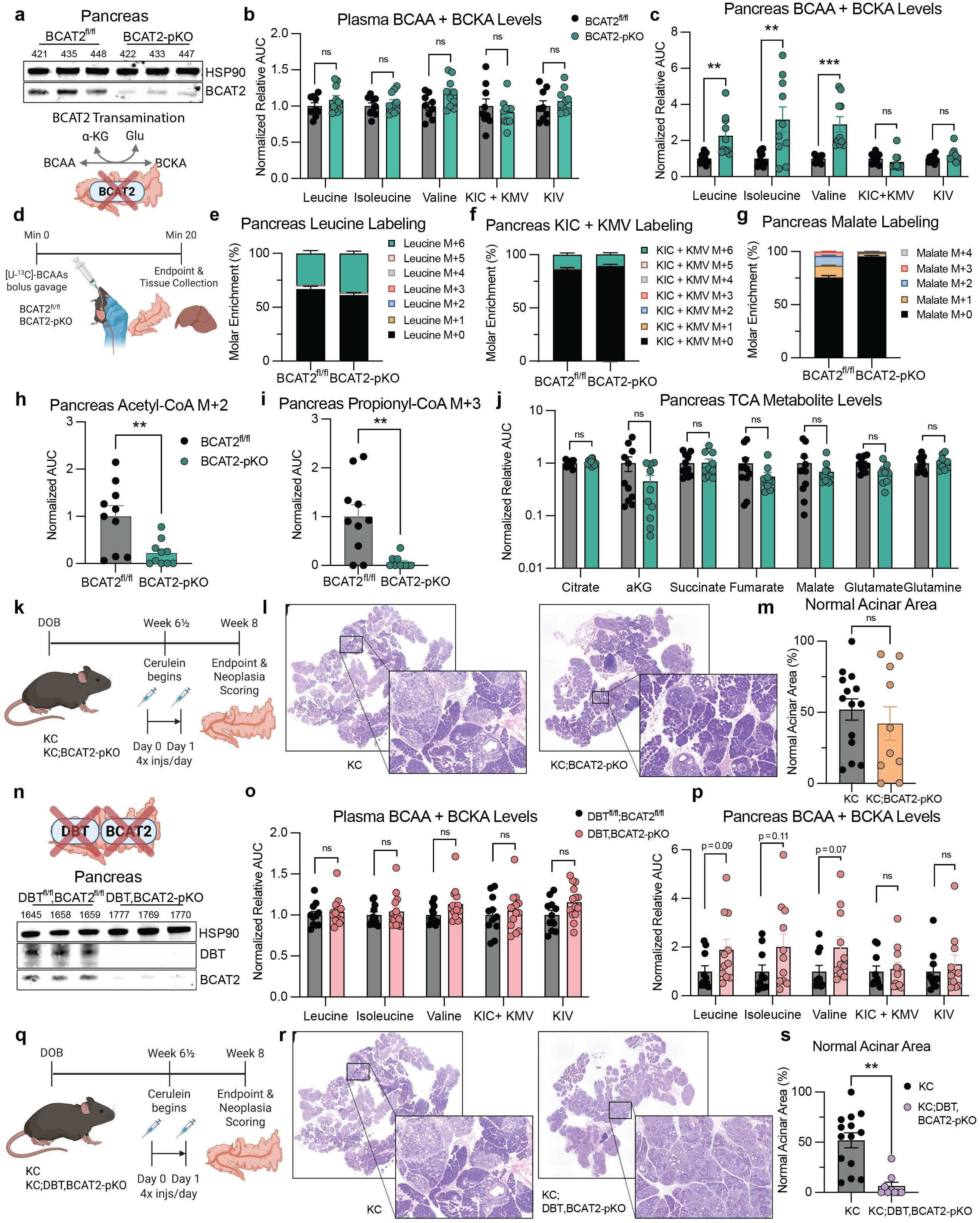
Neither metabolites upstream nor downstream of BCKDH explain the exacerbated PanIN formation phenotype in the absence of DBT. **a,** Western blot for BCAT2 protein in BCAT2^fl/fl^ and BCAT2-pKO pancreata collected from 8-10-week-old mice (n=3 BCAT2^fl/fl^ and n=3 BCAT2-pKO; HSP90 is the loading control). **b**, LC-MS measurements of relative plasma BCAAs and BCKAs of 5-hour fasted animals (n=9 BCAT2^fl/fl^ and n=10 BCAT2-pKO). **c**, LC-MS relative measurements of pancreas BCAAs and BCKAs in fed animals (n=11 BCAT2^fl/fl^ and n=10 BCAT2-pKO). **d**, Schematic diagram of bolus [U-^13^C]-BCAA gavage. **e-g**, MPE of **e**, leucine **f**, KIC + KMV, and **g**, malate in mouse pancreata 20 mins after bolus gavage (n=10 BCAT2^fl/fl^ and n=10 BCAT2-pKO). **h,i**, Acetyl-CoA M+2 and propionyl-CoA M+3 labeling measured in animal pancreata and liver 20 mins after bolus gavage of [U-^13^C]-BCAAs (n=10 BCAT2^fl/fl^ and n=10 BCAT2-pKO). **j**, LC-MS relative measurements of pancreatic TCA metabolites (n=11 BCAT2^fl/fl^ and n=10 BCAT2-pKO). **k**, Schematic and timeline for KC;BCAT2-pKO tumor initiation assays. **l**, Representative HCE images of KC;BCAT2-pKO animals after induction of pancreatitis at 6 ½ weeks and collected at an 8-week timepoint. **m**, Normal acinar area was plotted relative to total pancreas area (n=14 KC and n=10 KC;BCAT2-pKO). **n**, Western blot of DBT and BCAT2 proteins in animal pancreata at 8-10 weeks old (n=3 DBT^fl/fl^;BCAT2^fl/fl^ and n=3 DBT,BCAT2-pKO; HSP90 is the loading control). **o**, LC-MS relative measurements of plasma BCAAs and BCKAs of 5-hour fasted animals (n=11 DBT^fl/fl^;BCAT2^fl/fl^ and n=14 DBT,BCAT2-pKO). **p**, LC-MS relative measurements of pancreas BCAAs and BCKAs in fed animals (n=10 DBT^fl/fl^;BCAT2^fl/fl^ and n=11 DBT,BCAT2-pKO. **q**, Schematic and timeline for KC;DBT,BCAT2-pKO tumor initiation assays. **r**, Representative HCE images of KC;DBT,BCAT2-pKO animals after induction of pancreatitis at 6 ½ weeks and collected at an 8-week timepoint. **s**, Normal acinar area relative to total pancreas area at endpoint (n=14 KC and n=9 KC;DBT,BCAT2-pKO). Data are presented as mean ± s.e.m. Representative images depict animals closest to the mean of each group. KC historical data presented in Figure 3g re-graphed for comparison purposes for panels **m** and **s**. For comparing two groups, a two-tailed Student’s t-test was used for each individual metabolite plotted, with significance defined as **p<0.01 and ***p<0.001. **c**, Leucine *P*=0.0018; Isoleucine *P*=0.0046; Valine *P*=0.0002. **h**, Acetyl-CoA M+2 *P*=0.0044. **i**, Propionyl-CoA M+3 *P*=0.0014.

We first tested the metabolic effects of BCAT2 deficiency. BCAT2-pKO animals did not have overt changes in weight or tissue architecture (Extended Data Fig. 4b,c). No differences were observed in circulating plasma BCAA or BCKAs (Fig. 4b) and consistent with the transamination activity of BCAT2 within the pancreas, BCAA but not BCKA levels, were elevated (Fig. 4c). Interestingly, [U-^13^C]-BCAA gavage revealed labeling of both BCAAs and BCKAs (Fig. 4d-f) within the pancreas although BCKA pool size was reduced in the gavage setting in BCAT2-pKO animals (Extended Data Fig. 4d,e). [U-^13^C]-BCAA labeling of BCKAs was also recapitulated in BCAT2 deficient isolated acinar cells, suggesting that BCAT1 is likely active (Extended Data Fig. 4g-i). Importantly, however, BCAT2 deficiency ablated labeling by [U-^13^C]-BCAA of acetyl-CoA, propionyl-CoA, succinyl-CoA, and TCA cycle intermediates, with minimal change to TCA intermediate abundance, similar to that observed with DBT deficiency (Fig. 4g-j and Extended Data Fig. 4f-n). These data demonstrate that the BCKAs destined for oxidization in the pancreas are generated uniquely by BCAT2 (and not BCAT1), consistent with the existence of previously described BCAT2/BCKDH metabolons^35^. Thus, loss of either DBT or BCAT2 is sufficient to ablate pancreas BCAA catabolism.

To examine PanIN formation, we next generated KC;BCAT2-pKO animals and treated them with cerulein (Fig. 4k), as we had done for KC;DBT-pKO animals. Surprisingly, however, KC;BCAT2-pKO animals did not phenocopy the KC;DBT-pKO mice but rather developed PanINs comparably to KC control animals (Fig. 4l,m), and no significant differences in animal weight or relative pancreas mass were observed (Extended Data Fig. 5a,b). Similarly, no differences in PanIN formation were observed between non-cerulein treated KC and KC;BCAT2-pKO animals, further corroborating the finding that BCAT2 deficiency, unlike DBT deficiency, does not accelerate PanIN formation (Ext. Fig. 5c-g), even though loss of either enzyme suppresses BCAA oxidation. These data demonstrate that the loss of mitochondrial BCAA catabolism in pancreas is not sufficient to explain the exacerbated tumor formation in KC;DBT-pKO mice.

**Figure 5.**
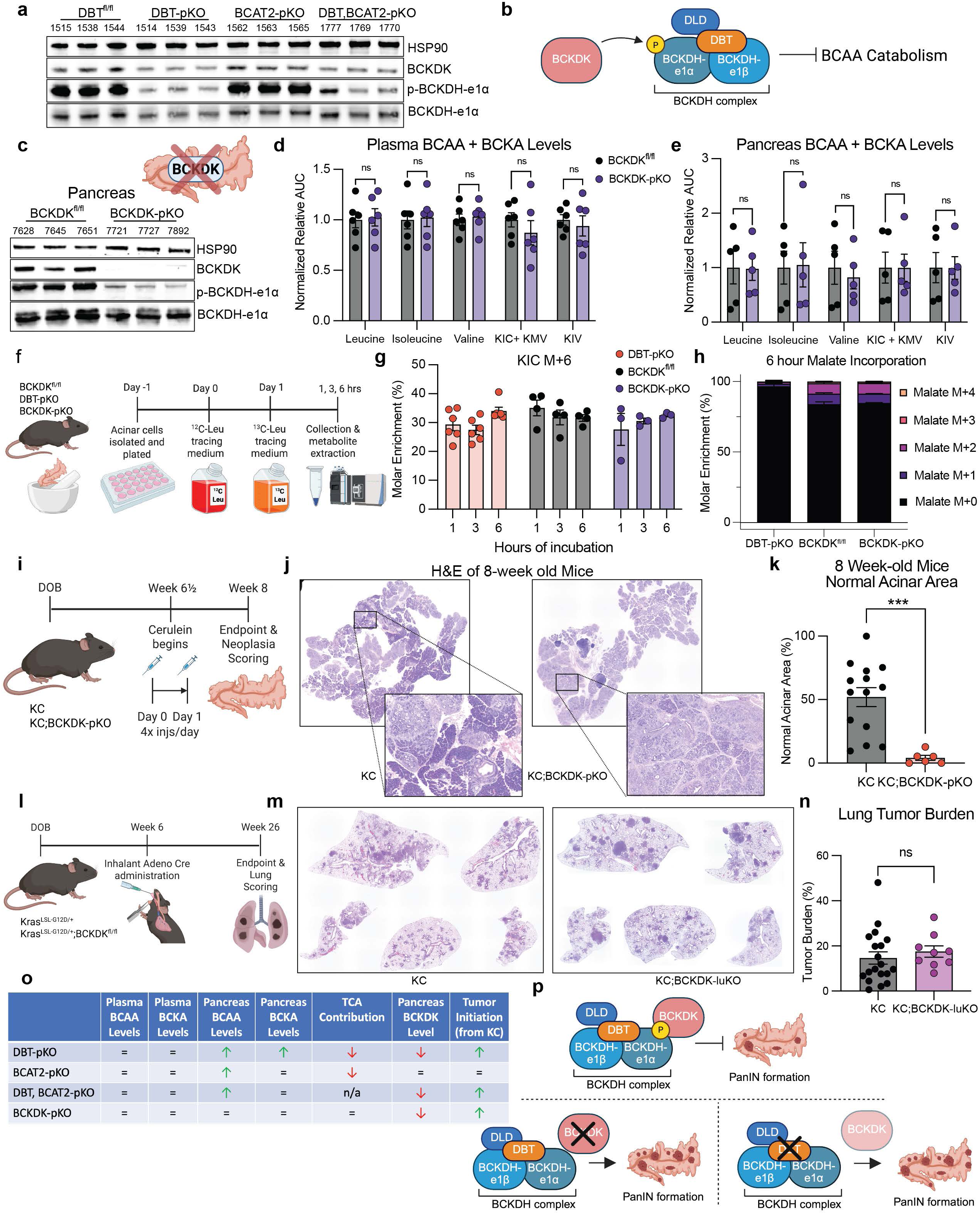
BCKDK abundance and activity are reduced in mice lacking pancreatic DBT and loss of pancreatic BCKDK increases PanIN formation. **a**, Western blots in pancreata from indicated genotypes of 8-10-week-old animals (n=3 per genotype; HSP90 is the loading control). **b**, Schematic diagram showing BCKDK phosphorylating BCKDH-e1α in the BCKDH complex which inhibits BCAA catabolism. **c**, Western blots in pancreata from 8-10-week-old animals (n=3 BCKDK^fl/fl^ and n=3 BCKDK-pKO; HSP90 is the loading control). **d**, LC-MS relative measurements of plasma BCAAs and BCKAs of 5-hour fasted animals (n=6 BCKDK^fl/fl^ and n=6 BCKDK-pKO). **e**, LC-MS relative measurements of pancreas BCAAs and BCKAs in fed animals (n=6 BCKDK^fl/fl^ and n=6 BCKDK-pKO). **f**, Schematic and timeline for *ex vivo* acinar cell [U-^13^C]-leucine tracing assay using freshly isolated acinar cells from indicated genotypes. **g**, KIC M+6 MPE shown after 1, 3, and 6 hours of incubation with [U-^13^C]-leucine (n=6 DBT-pKO, n=4 BCKCK^fl/fl^, and n=3 BCKDK-pKO). **h**, Malate MPE shown after 6 hours of incubation (n=6 DBT-pKO, n=4 BCKCK^fl/fl^, and n=3 BCKDK-pKO). **i**, Schematic and timeline for KC;BCKDK-pKO tumor initiation assays. **j**, Representative HCE images of KC;BCKDK-pKO animals after induction of pancreatitis at 6 ½ weeks and collected at an 8-week timepoint. **k**, Normal acinar area relative to total pancreas area (n=14 KC and n=6 KC;BCKDK-pKO). **l**, Timeline for tumor initiation assays for inhalant Ad:CMV-Cre Kras^G12D^ (KC) initiation model of lung adenocarcinoma for BCKDK-pKO animals. **m**, Representative HCE images of lung lobes 20 weeks after inhalant Cre delivery. **n**, Lung tumor burden is the percent of affected lung relative to total lung area (n=19 KC and n=9 KC:BCKDK-luKO). **o**, Table summarizing phenotypic observations across animal models. **p**, Model depicting BCKDK observations and PanIN formation rates. Data are presented as mean ± s.e.m. Representative images depict animals closest to the mean of each group as determined in panel k. KC historical data presented in Fig. 3c re-graphed for comparison purposes for pancreas and Fig. 3l re-graphed for lung comparisons. For comparing two groups, a two-tailed Student’s t-test was used for each individual metabolite plotted, with significance defined as ***p<0.001. **k**, Healthy acinar area *P*=0.0007

### Elevated BCKAs are also not sufficient to exacerbate PanIN formation

One difference between the DBT and BCAT2 deficient models is the accumulation of BCKAs, which is observed only in DBT-pKO mice. BCKAs can act as toxic metabolites and have been reported in patients with BKCDH deficiency, like those with Maple Syrup Urine Disease^36,37^ and their elevation can promote oxidative stress^38,39^, a known promoter of ADM and PanIN formation^40,41^. We thus hypothesized that the elevated BCKA levels in KC;DBT-pKO mice promote PanIN formation. To test this, we generated mice lacking both pancreatic DBT and BCAT2 (DBT,BCAT2-pKO) (Fig. 4n and Extended Data Fig. 6a). DBT,BCAT2-pKO mice are viable, grow similarly to littermate controls (Extended Data Fig. 6b) and pancreata show no overt differences by HCE staining or in expression of genes associated with pancreas function (Extended Data Fig. 6c,d). Importantly, the elevation in pancreas BCKAs observed with DBT deficiency alone was indeed reversed in the DBT,BCAT2-pKO mice, while similar to the single KO models, circulating levels of BCAAs and BCKAs were not altered (Fig. 4o,p). TCA metabolite abundance in DBT,BCAT2-pKO pancreata also showed no differences compared to controls (Extended Data Fig. 6e). We next generated KC;DBT,BCAT2-pKO mice and treated them with cerulein, anticipating that the resolution of BCKA elevation in these animals would protect from the accelerated PanIN formation observed in KC;DBT-pKO mice (Fig. 4q). Instead, however, KC;DBT,BCAT2-pKO animals phenocopied KC;DBT-pKO mice, with almost no remaining healthy acinar area (Fig. 4r,s) and significantly heavier pancreata at endpoint (Extended Data Fig. 7a,b). Increased PanIN formation in DBT,BCAT2-pKO mice was similarly seen in the absence of cerulein treatment (Extended Data Fig. 7c-j). These data demonstrate that elevated BCKAs are unlikely to be responsible for the increased PanIN formation observed with DBT deficiency.

### BCKDK protein expression and activity is impaired in mice lacking pancreatic DBT

Given the unexpected findings that both the metabolic pathways upstream and downstream of DBT are unlikely to account for the PanIN phenotype observed with DBT deficiency, we considered that perturbation of the BCKDH complex might have unanticipated effects on either other dehydrogenase complexes or on other BCKDH complex members. The BCKDH complex is in a family of related dehydrogenase complexes with pyruvate dehydrogenase (PDH) and oxoglutarate dehydrogenase (OGDH), all of which include the common subunit DLD. The levels of PDH-e1α, OGDH, and DLD did not change in the DBT-pKO mice (Extended Data Fig. 2b and Extended Data Fig. 8a). On the other hand, we noted that in DBT-pKO animal, BCKDH-e1α protein levels modestly decreased, and phosphorylation of BCKDH-e1α was drastically suppressed (Extended Data Fig. 8a and Fig. 5a). The kinase responsible for phosphorylating BCKDH-e1α is BCKDK (Fig. 1a and Fig. 5b). BCKDK protein abundance was also suppressed in the DBT-pKO and DBT,BCAT2-pKO pancreata, but not BCAT2-pKO mice, paralleling the accelerated tumor initiation phenotype in these animals (Fig. 5a). PPM1K, the phosphatase which removes phosphorylation on BCKDH-e1α was not changed (Extended Data Fig. 8a). We did not observe differences in p-PDH-e1α (Extended Data Fig. 8a), which is phosphorylated by different enzymes. Thus, loss of DBT but not BCAT2 results in suppressed BCKDK abundance in the pancreas, correlating with the enhanced tumor initiation phenotype.

### Loss of BCKDK in the pancreas leads to increased PanIN formation

Our data led us to suspect that the loss of BCKDK function might lead to accelerated PanIN formation, independently of its effect on BCAA oxidation. To test this, we leveraged *Bckdk* floxed animals^42^ to produce mice lacking BCKDK in the pancreas (BCKDK-pKO mice) (Fig. 5c). These animals showed no difference in body weight, in tissue architecture and integrity, in pancreas-function related gene expression, or in glucose tolerance (Extended Data Fig. 8b-e). No changes were observed in pancreas or circulating levels of BCAAs and BCKAs or in abundance of TCA cycle intermediates in pancreas (Fig. 5d,e and Extended Data Fig. 8f). Notably, BCKDK deficiency also did not alter BCAA-dependent labeling of TCA cycle intermediates in an *ex vivo* acinar cell model (Fig. 5f-h and Extended Data Fig. 8g,h), suggesting that since pancreas BCAA catabolism occurs at a high rate^26,27^, catabolism of BCAAs in the pancreas is not normally inhibited by BCKDK. Nevertheless, KC;BCKDK-pKO mice treated with cerulein exhibited increased PanIN development, heavier pancreata, and reduced normal acinar area (Fig. 5i-k and Extended Data Fig. 9a,b), similar to that observed in KC;DBT-pKO mice. Animals not treated with cerulein also showed less normal acinar area at both 8 and 12-weeks old at endpoint (Ext. Fig. 9c-k). Also similar to that observed with DBT deficiency, lung tumor formation was not altered by BCKDK loss (Fig. 5l-n and Extended Data Fig. 9l). Altogether, these data indicate that although BCKDK exerts minimal effect to restrain BCAA catabolism in the healthy pancreas, loss of pancreas BCKDK unexpectedly accelerates KRAS-driven PanIN formation (Fig. 5o,p), suggesting that BCKDK may exert these effects through mechanisms independent of BCAA metabolism.

### Increasing BCAA catabolism pharmacologically decreases circulating BCAA levels without further accelerating pancreatic BCAA catabolism

The data above demonstrate a key role for the pancreatic BCKDH complex and BCKDK, but not BCAA oxidation *per se*, in PanIN formation. The data do not, however, address the potential effects of targeting systemic BCKDK or BCAA catabolism. Notably, prior work has shown that limiting systemic BCAA availability through the diet has a modest but significant effect in restraining tumor formation^23^, and systemic treatment with the BCKDK inhibitor 3,6-dichlorobenzo[b]thiophene-2-carboxylic acid (BT2)^43,44^ lowers circulating levels of BCAAs and BCKAs^26,42,45^ and may reduce the growth of PDA tumors implanted subcutaneously (though in nude mice with no intact immune system)^21^. Interestingly, BT2 administration reduces p-BCKDH-e1α across multiple tissues, but not the pancreas^26^, suggesting that BT2 might inhibit systemic BCKDK without substantial effect on pancreas BCKDK. We therefore tested the effect of BT2 in the KC model. To start, we provided 4-week-old KC animals BT2-containing chow or an isocaloric control chow (Fig. 6a). BT2 chow-fed animals grew at similar rates as control (Extended Data Fig. 10a). Fasted plasma BT2 levels after 1 week on diet confirmed the presence of BT2 in circulation (Extended Data Fig. 10b) and target engagement was confirmed by modest decreases in circulating levels of BCAAs and striking decreases in BCKAs (Fig. 6b), consistent with prior observations^26,42,45^. BT2 was detected in tissues including the pancreas and liver (Extended Data Fig. 10c), but no differences were seen in pancreas BCAAs, BCKAs, or p-BCKDH-e1α levels in BT2-fed animals (Fig. 6c,d and Extended Data Fig. 10d), as previously seen in WT animals^26^. As expected, BT2 reduced p-BCKDH-e1a in heart and liver (Fig. 6d and Extended Data Fig. 10e), concordant with prior work^26^. No change in liver pool size for BCAAs, BCKAs, or TCA intermediates was observed (Extended Data Fig. 10e-g). Thus, BT2 promotes systemic, but not pancreatic BCAA catabolism.

**Figure 6.**
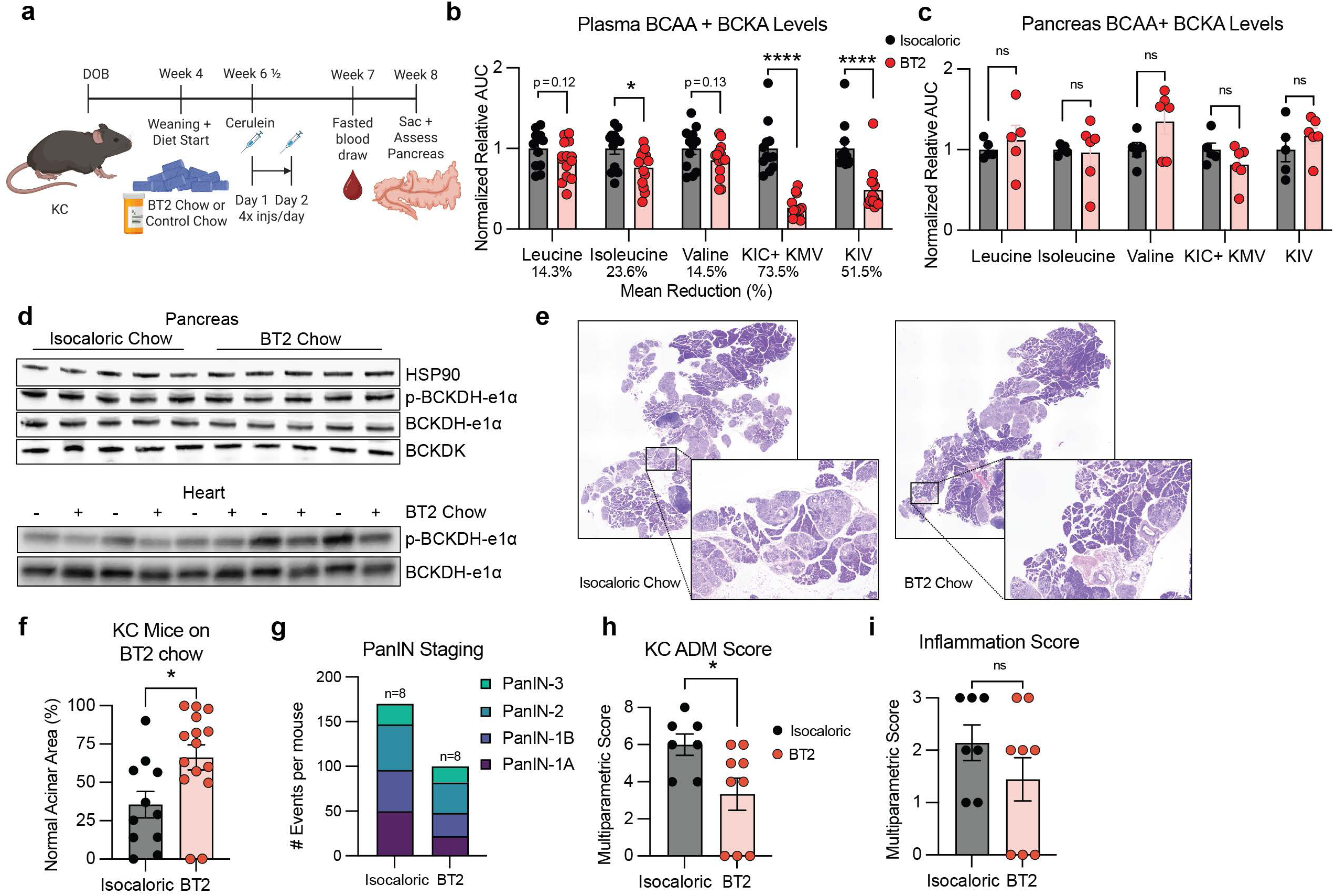
Systemic BCKDK inhibition, in contrast to pancreas BCKDK loss, reduces PanIN formation. **a**, Schematic and timeline for KC animals fed BT2 chow and collected at 8 weeks old after induction of pancreatitis with cerulein at 6 ½ weeks old as previously done. **b**, LC-MS relative measurements of plasma BCAAs and BCKAs after 3 weeks on BT2 chow and 5-hour fast (n=12 isocaloric chow and n=14 BT2 chow). **c**, LC-MS relative measurements of pancreas BCAAs and BCKAs at fed state after 3 weeks on diet (n=5 isocaloric chow and n=6 BT2 chow). **d**, Western for p-BCKDH-e1α, BCKDH-e1α, and BCKDK protein measured in pancreas and heart after 1 week on diet (n=5 per genotype; HSP90 is the loading control). Animals were harvested in fed state. **e**, Representative HCE images of KC BT2 and isocaloric fed animals at endpoint. **f**, Normal acinar area relative to total pancreas area (n=11 isocaloric chow and n=15 BT2 chow). **g**, Histopathological scoring of number and severity of PanIN lesions (n=8 isocaloric chow and n=8 BT2 chow; 8 animals from **f** closest to the mean within each group were chosen). **h,i**, Histopathological scoring of ADM and Inflammation as graded on a multiparametric scale. Scoring is broken down in Extended Data Fig. 3c,d. Data are presented as mean ± s.e.m. Representative images depict animals closest to the mean of each group. For comparing two groups, a two-tailed Student’s t-test was used for each individual metabolite plotted, with significance defined as *p<0.05 and ****p<0.0001. **b**, Leucine *P*=0.1217; Isoleucine *P*=0.0174; Valine *P*= 0.1293; KIC+ KMV *P*<0.0001; KIV *P*<0.0001. **f**, Normal acinar area *P*=0.0179. **h**, ADM Score *P*=0.0308.

Strikingly, the healthy acinar area was better preserved in pancreata of BT2-fed animals (Fig. 6e,f), with a slight decrease in pancreas mass (Ext. Fig. 10h). Histopathological scoring confirmed that BT2-fed animals had significantly fewer and lower grade PanINs, as well as a lower ADM score and a trend towards a reduced inflammation score (Fig. 6g-i). All together, these data demonstrate that (1) BCKDK in the pancreas is protective against PanIN formation; but that (2) pharmacologic inhibition of systemic BCKDK and promotion of systemic BCAA catabolism can be used to slow pancreatic cancer initiation. The latter is likely in part enabled because BT2 does not appear to inhibit BCKDK in the pancreas itself (and thus not inhibiting its protective effects).

## Discussion

In this work we identify contrasting effects of BCKDK inhibition systemically versus within the pancreas on PanIN formation. Suppression of BCKDK limited to the pancreas, whether directly by genetic deletion of *Bckdk* or indirectly by reduced protein abundance in the absence of DBT, accelerates PanIN formation, while in contrast, systemic inhibition of BCKDK, achieved pharmacologically with BT2, protects against PanIN formation. We discuss these two effects separately below.

Systemic treatment with BT2 does not appear to inhibit BCKDK in the pancreas itself (based on phosphorylation of BCKDH-e1α as a readout), indicating that the suppression of PanIN formation by systemic BT2 treatment is likely mediated on cells originating outside the pancreas. A key question arising from our work is thus what other cell types might be responsible for this strong suppression of PanIN formation. Inflammation plays an important role in PDA initiation, and immune cells could be impacted by BT2. BCAAs are known to modulate immunometabolism and immune cell function^46^, including regulating macrophage polarization^47^ and CD8^+^ T-cell effector function^48^. In CAR-T cells, BCKDK was shown to promote anti-cancer function^49^. It will be of interest in future work to test the impact of *Bckdk* deletion in macrophages or in T-cells on the initiation and progression of PanINs. Another possibility is that the lowering of systemic levels of BCAAs and BCKAs by BT2 drives the suppression of PanINs. BT2 promotes BCAA oxidation in muscle, brown adipose tissue, liver, and heart^26^, reducing BCAAs and BCKAs in circulation, which may in turn indirectly affect PanIN formation or growth. Studies with deletion of *Bckdk* in these tissues could address this possibility. Finally, it should be noted that BT2 could be acting through mechanisms other than BCKDK inhibition. BT2 has been described to have off-target effects, such as increasing tryptophan catabolism^50^ and modulating mitochondrial reactive oxygen species (ROS)^51^, and in some contexts BT2 treatment does not recapitulate phenotypes of BCKDK deficiency^26,42,45,50^. It will therefore be of interest to test other inhibitors of BCKDK, e.g. recently derived structurally altered molecules^52,53^.

In sharp contrast to the benefits of systemic inhibition of BCKDK, loss of BCKDK specifically in pancreatic cells promotes PanIN formation. Importantly, this effect is independent of the canonical role of BCKDK in regulation of BCAA catabolism, since loss of either DBT or BCKDK-which have opposing roles in BCAA metabolism-both result in accelerated PanIN formation. Our data thus strongly suggest the involvement of other BCKDK substrates. BCKDK has been proposed to target other substrates including ACLY and PDH^54,55^. For example, when PDH kinase enzymes are missing during development, BCKDK can phosphorylate PDH^55^, although we did not observe differences in p-PDH-e1α in our model (Extended Data Fig. 8a). BCKDK may also have additional substrates that are not yet described. Future work with phospho-proteomics may identify such substrates. Finally, it is also possible that BCKDK has non-enzymatic roles in suppressing PanIN formation. Testing this possibility would require generation of mice expressing kinase-dead BCKDK in the pancreas.

These findings also highlight an interesting example of unanticipated effects of gene deletion *in vivo,* i.e. the loss of BCKDK protein upon the deletion of DBT. Roth Flach *et al.* showed that BT2 binding to BCKDK promotes its dissociation from the BCKDH complex and its subsequent degradation^52^, an observation that let us to test the impact of DBT deletion on BCKDK abundance. In effect, DBT-pKO mice became DBT,BCKDK-pKO mice, phenotypically mimicking BCKDK-pKO mice. Consistent with these observations, BCKDK has previously been purified and found to physically to bind to DBT, and not BCKDH-e1α, in biochemical assays^56^.

Our work also highlights the distinct roles that BCAA catabolism plays in tumor initiation versus tumor growth and progression. Despite the minimal contribution of BCAAs to the TCA cycle in PDA (Fig. 1d,e), several studies have shown slowed cell proliferation and reduced tumor growth upon targeting of BCAA enzymes, including BCAT2^57^ and members of the BCKDH complex (BCKDH-e1α^57,58^ and DBT^59^). Our lab has found that BCAA-derived propionyl-CoA is important for histone propionylation and transcription in established PDA cells^59^. In the tumor initiation setting in this study, in contrast, we find that propionyl-CoA production is suppressed by both DBT and BCAT2 KO, which have differing effects on PanIN development, making it unlikely that BCAA-dependent histone propionylation is a contributor to the PanIN formation phenotypes observed with DBT and BCKDK deficiency. Distinct from these observations in tumor growth models, our current data support the notion that BCAA catabolism has little impact on tumor initiation, i.e. PanIN formation, because loss of BCAT2 blocked BCAA oxidation in the pancreas but had no effect on PanIN formation, while loss of DBT, which also blocked BCAA oxidation increases PanIN formation. Our work does contrast with a previous report of reduced PanIN formation and elevated plasma BCAAs in mice lacking pancreatic BCAT2^23^; these differences may reflect differences in mouse background or environmental factors. Regardless, both studies are consistent with the conclusion that the loss of BCAA catabolism does not underlie the accelerated PanIN formation phenotype that we observe in the DBT deficient mice.

In conclusion, we have shown diverging roles of modulating systemic versus pancreas-intrinsic BCKDK in PanIN development at the onset of PDA. We find that pancreatic BCKDK loss exacerbates PanIN development, but in contrast, systemic promotion of BCAA catabolism by inhibiting BCKDK is protective against PanIN development. BCKDK promotes tumor growth and metastasis in several other contexts^49,60–63^ and its inhibition sensitizes cancer cells to standard-of-care cancer treatments such as paclitaxel^64,65^ or doxorubicin^66^. Moreover, BT2 has been shown to be beneficial in preclinical models of a variety of metabolic disorders, including cardiovascular disease^45^, insulin resistance^26,42,67^, acute kidney injury^68^, ulcerative colitis^69^, sarcopenia^70^, Alzheimer’s disease^71^, and bone inflammation and erosion^72^. Several BT2 analogs have been developed^52^ and a more potent analog, whose structure was recently disclosed^53^, is now in clinical trials (NCT05654181). Systemic BCKDK inhibition thus represents a promising strategy to reduce carcinogenesis in individuals with high risk factors for PDA, such as chronic pancreatitis^9^ or diabetes^6,7^, with at least one clinical candidate with positive safety profile already available.

## Methods

### Murine Animal Models

All animal studies were approved by the University of Pennsylvania (UPenn) Institutional Animal Care and Use Committee (IACUC). Mice were housed on a normal light-dark cycle (light on from 7 a.m. to 7 p.m.) and fed standard rodent chow (LabDiet, 5010), except when special diets or conditions are indicated. The temperature range in the animal housing facility was 21–23°C. Animals weights were recorded weekly in the afternoon. Generally, plasma sampling was done in the afternoon (1 p.m. to 3 p.m.) for fasted values, animals were fasted for 5 hours (8 a.m. to 1 p.m.) prior to blood collection. For experiments, plasma was collected the week prior to endpoint unless specifically noted. Forelimb size was measured at endpoint with calipers measuring from elbow to the base of the palm.

Generation of murine models with pancreas-specific deletions all used the *P48-Cre* driver^73^. These animals were gifted from Celeste Simon and were originally obtained from the MMRC (000435-UNC). For all breeding schemes, Cre was maintained in the maternal offspring. *DBT^fl/fl^*mice generation has previously been reported^42^ and the generation of *P48-Cre;DBT^fl/fl^*(pancreas-specific DBT deletion, DBT-pKO) animals was done on C57Bl6/J background. Specifically, female *P48-Cre* animals were crossed to *DBT^fl/fl^* males to generate *P48-Cre;DBT^fl/+^* animals. These were crossed to *DBT^fl/fl^* animals to generate *P48-Cre;DBT^fl/fl^* animals. These *P48-Cre;DBT^fl/fl^* females were then finally crossed to male *DBT^fl/fl^* animals to generate DBT-pKO litters. *BCAT2^fl/fl^* animals were gifted from Susan Hutson and their generation and phenotyping has been previously described^74^. Similar to DBT-pKO animals, BCAT2-pKO animals were generated by crossing *P48-Cre;BCAT2^fl/fl^* females to the *BCAT2^fl/fl^*male animals. The generation of *P48-Cre;DBT^fl/fl^;BCAT2^fl/fl^* (DBT, BCAT2-pKO) animals was similar to previous models where *P48-Cre;DBT^fl/fl^;BCAT2^fl/fl^* females were crossed to *DBT^fl/fl^;BCAT2^fl/fl^*male animals. To generate *DBT^fl/fl^;BCAT2^fl/fl^*animals, *DBT^fl/fl^*were crossed to *BCAT2^fl/fl^* animals to generate *DBT^fl/+;^BCAT2^fl/+^* animals. These heterozygous animals were then crossed to one another to generate *DBT^fl/fl^;BCAT2^fl/fl^* animals. *BCKDK^fl/fl^* animal generation has been previously described^42^ and these were crossed to *P48-Cre* animals, similarly to breeding schemes mentioned above.

For all PDA initiation experiments, we crossed our baseline pancreas-deletion animals to Kras^LSL-G12D/+^ animals, which have constitutively active Kras signaling, have been extensively studied and originally generated for lung cancer studies^75^. For example, to generate (*P48-Cre;Kras^LSL^-^G12D/+^,DBT^fl/fl^*) animals, we crossed the *Kras^LSL-G12D/+^*animals to *DBT^fl/fl^*to generate *Kras^LSL-G12D/+^;DBT^fl/+^*animals. These were then crossed to *DBT^fl/fl^*animals to generate *Kras^LSL-G12D/+^,DBT^fl/fl^* animals. Female *Kras^LSL-G12D/+^,DBT^fl/fl^* animals were then crossed to *P48-Cre;DBT^fl/fl^* animals to finally generate the *P48-Cre;Kras^LSL-G12D/+^,DBT^fl/fl^* animals which we refer to as KC;DBT-pKO animals. Crossing the *Kras^LSL-G12D/+^* mutation to other pancreas-specific models was done similarly to generate the KC;BCAT2-pKO, KC;DBT,BCAT2-pKO, and KC;BCKDK-pKO animals.

For the orthotopic tumor exogenous BCAA water assay and the effect of BT2 on organ BCKDH expression, male C57Bl6/J WT animals were purchased at 6 weeks old (Jackson Labs #000664).

Genotyping for all breeding was confirmed via Transnetyx (Cordova, USA) by sending an ear tag or tail snip per animal.

### Mammalian Cell Culture

Immortalized PDA cells were cultured in DMEM (Gibco 11965-084) with 10% Calf Serum (HyClone SH30072.03), 1% PenStrep (Invitrogen 15140122), and 1% Glutamax (Gibco 35050061). Cells were maintained in 21% O_2_, 5% CO_2_, and 37 °C standard culturing conditions unless otherwise noted. Cells were passaged using 0.05% trypsin (Invitrogen 25300054) to detach adherent cells from the plate.

### *Ex vivo* Acinar Cell Isolation

Mice were euthanized and pancreata harvested the morning of isolation as previously described^27^. In short, pancreata were isolated from the animal and transferred to a sterile biosafety cabinet in ice-cold PBS (Corning 21-031-M). In the hood, the pancreata were minced into small pieces (∼1 mm diameter) using sterile scalpels. These small pieces were enzymatically dissociated with 1 mg/mL Collagenase D (Millipore Roche 11088866001) for 45 minutes while rotating in a 37 °C incubator. After dissociation, the cells undergo a series of 5% CS HBSS (Gibco 14175) washes and centrifugal pelleting before they are passed through a 500 uM and then subsequently a 100 uM filter. Once in a single-cell suspension, cells were pelleted and resuspended in 10% CS Waymouth’s medium (Gibco SMB-7521) with 0.1mg/mL trypsin soybean inhibitor (Gibco 17075-029). These cells were then seeded into ultra-low-adhesion 24-well plates (Costar 3473) and cultured in standard cell conditions.

### ^13^C Isotope Tracing

Adherent immortalized PDA cells were plated in 6-well plates in triplicate at 100k cells/well. Cells were left to adhere overnight and the next day, media was aspirated off and cells were washed with PBS before basal tracing media with supplements was provided. Tracing medium is a DMEM-based medium without glucose, glutamine, and BCAAs (US Biological D9800-36). ^12^C glucose (25 mM), ^12^C glutamine (2 mM), ^12^C leucine (0.8 mM), ^12^C isoleucine (0.8 mM), ^12^C valine (0.8 mM) were added to base media along with 10% dialyzed fetal bovine serum (Gemini Bio 100-108) for 24 hours to allow for cells to acclimate to the new media. The next morning, the basal tracing medium was refreshed and 2 hours later, cells were washed 2x with dPBS before tracing media with ^13^C-metabolites was added. Control wells were provided with ^12^C tracing media, while ^13^C metabolite wells were provided with all ^12^C metabolites mentioned above except for the metabolite of interest (ie. [U-^13^C]-glutamine labeled cells receive ^12^C glucose, ^12^C BCAAs, but ^13^C glutamine. Concentrations of ^13^C metabolites were equal to ^12^C so that cells do not experience any change in metabolite abundance during the tracing period. Cells were incubated with labelled substrates for 6 hours before they were harvested for extraction to be run on GC-MS/LC-MS. Unlabeled control wells were used to correct labeling for natural abundance of isotopologues.

Freshly isolated pancreatic acinar cells were cultured in 10% CS Waymouth’s Medium and put into ultra-low attachment 24-well plates overnight. The next day, cells were collected from wells and spun down in a 4°C table-top centrifuge at 400 x g for 4 minutes. Cells were then re-suspended in basal tracing medium as mentioned above with ^12^C metabolites for 24 hours to allow for cells to acclimate to the new medium. The next day, fresh ^12^C tracing medium was refreshed for 2 hours. Cells were then washed 2x with PBS before tracing medium with ^13^C label was introduced as described above. Cells were incubated for 6 hours before they were harvested and extracted.

### Metabolite Extraction for GC-MS

#### Cells

Cells were collected at experimental end point and extracted using 80% MeOH (mass-spec grade diluted in water) on ice. For adherent cells, cells were washed with ice cold PBS 2x prior to adding extraction solution. Once added, cells were scraped using a cell lifter (on ice. For anchorage-independent cells, cells were pelleted and extraction buffer was directly added to the pelleted cells on ice. Extraction solution was spun down in a 4 °C table-top centrifuge and the supernatant was saved. The pellet was saved at -80 °C. For extraction normalization, an internal standard (norvaline) was added to each sample pellet to account for sample loss during prep. Samples were then vortexed for 60 seconds prior to centrifugation at 4 °C at max speed for 15 minutes. The supernatant was saved and was then spun down overnight to dry using a Speedvac (Thermo Scientific SPD130DLX).

Once dried, metabolite pellets were resuspended in 30 uL pyridine containing methoxyamine HCl (Sigma 226904; 10 ug/uL). These samples were then vortex for 10 seconds each, twice. These were then heated at 70 °C for 15 minutes on a heating block. 70 uL MTBSTFA (Sigma 394882) was added to these samples for derivatization and heated further at 70 °C for 60 minutes after a quick 5 second vortex. Samples were then spun down at full speed for 20 minutes at 4 °C, and 40uL of the supernatant was transferred to glass vials with glass inserts and saved at 4 °C until they were run on the GC-MS (Agilent 8890/5977).

### GC-MS Analysis for Polar Metabolites

One microliter of each sample was injected by the automatic liquid sampler into the GC. The GC method used splitless injection mode with 1.2 ml min^−1^ helium as a carrier gas moving each sample through a 30 m × 250 μm × 0.25 μm HP-5ms Ultra Inert GC Column (Agilent 19091S-433UI). The inlet temperature was 250 °C and the oven temperature was initially 100 °C. After 3 min, the oven temperature was increased by 4 °C per minute to 230 °C then by 20 °C per minute to 300 °C and was held for 5 min. The transfer line temperature was 250 °C, the MSD source temperature was 230 °C and the MSD quadrupole temperature was 150 °C. The method included a 6-min solvent delay, after which the MSD operated in electron ionization mode and scan mode with a mass range of 50–550 atomic mass units at 2.9 scans per second. The Agilent MassHunter Quantitative Analysis software was used to quantify chromatograms obtained from the above method. The relative abundance of each metabolite was normalized to an internal standard (norvaline) for each sample. Normal isotopic distribution was corrected ^13^C natural abundance for using FluxFix^76^ and isotopologue enrichment data was shown as molar polar enrichment and total pool size (stacked isotopologues of each metabolite of interest).

### Metabolite Extraction for LC-MS

#### Tissues

Animal tissues were collected and immediately freeze-clamped in aluminum foil in liquid nitrogen. Once frozen, all tissues were kept at -80 °C until time of extraction. Tissues were kept frozen on dry ice while we mechanically broke up the tissue into small pieces (∼1-2 mg). Small pieces from regions throughout the tissue were pooled and weighed while remaining frozen. Roughly 20-30 mg of tissue was saved for extraction with exact weights recorded for biological normalization purposes. 1 mL of ice-cold extraction buffer (+ internal standard) was added to the tissues. This internal standard was either U-^13^C-propionate or U-^13^C-isoleucine, depending on the experiment. Samples were then homogenized with a benchtop lyser (SciLogex OS20-S) at 1600 mhZ for approximately 15 seconds. Samples were then immediately centrifuged at max speed for 20 mins at 4 °C and the supernatant was saved at -80 °C.

#### Plasma

Animal blood was collected via-tail bleeds either in a fed state (∼1-3 PM) or after a 5-hour fast (8AM-1 PM) using EDTA-containing plasma collection tubes (Sarstedt 16-443-100). These were spun down at 2,500 RPM for 15 minutes at 4 °C before the supernatant plasma was saved and frozen at -80 °C. For extraction, 3 uL of plasma was extracted in 260 uL extraction buffer (+ internal standard) on ice for 10 minutes. Samples were then immediately centrifuged at 13,300 g for 20 mins at 4 °C and supernatant was saved at -80 °C.

Regardless of biological input, extracted supernatant was dried down using a SpeedVac. Once dried, metabolite pellets were resuspended in ACN:H_2_O in a ratio of 60:40 and incubated on ice for 15 minutes. Every 5 minutes, samples were vortexed for 10 seconds. After 15 minutes, samples were spun down and 40 uL of the supernatant was transferred to glass vials with glass inserts and saved at 4 °C until they were run on the LC-MS (Thermo Fisher Q Exactive quadropole-orbitrap mass spectrometer).

### LC-MS Analysis for Polar Metabolites

A quadrupole-orbitrap mass spectrometer (Q Exactive, Thermo Fisher Scientific) operating in negative ion mode was coupled to hydrophilic interaction chromatography on a Vanquish UHPLC System (Thermo Fisher Scientific) via electrospray ionization and used to scan from *m*/*z* 65–425 at 1 Hz and 140,000 resolution. LC separation was on a ACQUITY Premier BEH Amide VanGuard FIT Column, 1.7 μm, 2.1 mm × 100 mm (Waters) using a gradient of solvent A (20 mM ammonium acetate, 20 mM Ammonium hydroxide in 95:5 H_2_O:ACN, pH 9.45) and solvent B (100% ACN). Flow rate was 300 μL/min. The LC gradient was: 0 min, 95% B; 9 min, 40%; 11 min, 40%; 11.1 min, 95%; 20 min, 95%. Autosampler was set at 4 °C and injection volume was 3 μL. Metabolite values are shown after being normalized to the internal standard for each sample and also after normalizing to biological input. For tissues, this was the weight of the sample weighed out for extraction. Data analysis was done using El-MAVEN software^77^. Normal isotopic distribution were corrected for ^13^C natural abundance using FluxFix^76^ and isotopologue enrichment data were shown as molar polar enrichment and total pool size (stacked isotopologues of each metabolite of interest).

### U-^13^C-BCAA Gavage for Polar Metabolites and CoA Tracing

8–10-week-old animals were placed in individual containers in the afternoon. The U-^13^C-BCAA gavage solution was prepared in saline with a supraphysiological concentration of BCAAs (82.5 mM Leucine, 37.5 mM Isoleucine, and 60 mM Valine). This solution was filter-sterilized and animals were given an oral gavage (8 uL/g animal mass) which was 180 mg/kg. 20 minutes later, animals are quickly euthanized via cervical dislocation in accordance to our IACUC protocol. Tissues of interest (pancreas, liver) were excised and quickly freeze-clamped in aluminum foil in LN_2_. Cardiac puncture was also performed to collect plasma at endpoint. Tissues were saved at -80 °C until time of extraction. At time of extraction, tissues were kept frozen as described earlier and 20-30 mg were weighed out and recorded. These samples were then kept at -80 °C until they were extracted either using 40:40:20 ACN:MeOH:H_2_O for polar metabolites as described earlier or 10% TCA for CoA species.

### Metabolite Extraction for Acyl-CoAs

Acyl-CoAs were analyzed by liquid chromatography-high-resolution mass spectrometry (LC-HRMS) as previously described^78^. 50 µL of short-chain acyl-CoA ISTD was added and then cell suspensions were sonicated with 5 x 0.5-second pulses at 50% intensity (Fisherbrand™ Sonic Dismembrator Model 120 with Qsonica CL-18 sonicator probe). Lysates were centrifuged at 17,000 x g for 10 minutes at 4 °C and clarified lysates were transferred to a deep-well 96-well plate for loading in a Tomtec Quadra4 liquid handling workstation. On the liquid handling workstation, lysates were applied to an Oasis HLB 96-well elution plate (30 mg of sorbent per well) pre-conditioned and equilibrated with 1 mL of methanol and 1 mL of water, respectively. After de-salting with 1 mL of water, acetyl-CoA was eluted into a deep-well 96-well plate using 1 mL of 25 mM ammonium acetate in methanol. Eluent was evaporated dried under nitrogen gas. The dried LC-HRMS samples were resuspended in 50 µL of 5% (w/v) sulfosalicylic acid in water. 5 µL injections of each sample were analyzed via LC-HRMS. Acetyl-CoA, propionyl-CoA, and succinyl-CoA lithium salts as well as 5-sulfosalicylic acid were from Sigma-Aldrich. Optima® LC/MS grade acetonitrile (ACN), formic acid, methanol, and water were purchased from Fisher Scientific. Oasis® HLB 96-well elution plates (30 mg of sorbent) were purchased from Waters (P/N: WAT058951). Short-chain acyl-CoA internal standard (ISTD) was generated in yeast as previously described^79^.

### LC-MS Analysis for Acyl-CoAs

Samples were analyzed using a Vanquish Duo ultra-high performance liquid chromatograph coupled with a Q Exactive Plus mass spectrometer (Thermo Scientific) as previously described^78^. A modified gradient using solvent A (5mM ammonium acetate in water), solvent B (5 mM ammonium acetate in 95:5 (v:v) acetonitrile: water) and solvent C (0.1% (v/v) formic acid in 80:20 (v:v) acetonitrile: water). Data were acquired using XCalibur 4.0 (Thermo Scientific), analyzed using Tracefinder 5.1 (Thermo Scientific), and corrected for normal isotopic distribution using FluxFix^76^.

### Histology

Animal tissues were placed in formalin in the well of a 6-well plate for overnight fixation. Pancreata were first spread onto filter paper (Bio-Rad) cut into a 1” diameter circle and placed in the bottom of the 6-well plate to prevent tissue from curling up. After overnight fixation, tissues were placed into cassettes and dehydrated through a 50% ethanol for 1 hour and a subsequent 70% ethanol wash. Once in 70% ethanol, tissues were submitted to the University of Pennsylvania’s Center for Molecular Studies in Digestive and Liver Diseases for paraffin-embedding, slide-mounting, and hematoxylin and eosin (HCE) staining. HCE slides were imaged at 2X magnification and stitched together using a Keyence BZ-X710. Merged images were then opened in Fiji^80^ for assessment and quantification.

Pancreata of KC animals were first quantified for total area and then quantification for non-healthy, aberrant pancreas tissue was done using the FIJI measure tool to calculate areas I had outlined for quantification. This aberrant, non-healthy pancreas area includes pancreatic intraepithelial neoplasia and early-stage PDA. After measuring the total area of the pancreas and that of the non-healthy tissue, a percentage of the healthy acinar area (non-aberrant) was calculated. This is how PDA initiation was assessed in our animal models. This area calculation was similarly done for the lung tumor burden (tumor area/total area) and Ki-67^+^ staining (Ki-67^+^ nuclei/total nuclei per slide).

### Glucose Tolerance Tests (GTTs)

Animals were individually housed on ALPHA-dri bedding (Shepard Specialty Papers) and fasted in the morning for 5 hours starting at 7 AM. A time 0 blood sample was acquired via tail snip. After blood samples were collected for all animals, mice were orally gavaged with 20% dextrose solution (diluted in saline) at 2 g/kg per animal using a 2 mL syringe with a plastic feeding tube (Instech FTP-20-38). Animals were staggered 3 minutes apart to allow time for sampling. At 20 minutes post injection, blood was sampled again and collected to assess individual insulin response to the dextrose gavage. At 0, 20, 40, 60, and 120 minutes post injection, a glucometer (Contour Next) was used to measure blood glucose values. Males and females were run on separate days for experiments.

### Insulin enzyme-linked immunosorbent assays (ELISAs)

Insulin concentrations were measured from plasma taken at 0 and 20 minutes post dextrose gavage during the GTT. The “wide range assay” used was described in the Ultra Sensitive Insulin ELISA Kit (Crystal Chem #90080).

### RNA Extraction and qPCR Analysis

Animals were euthanized in the afternoon and immediately the tissue of interest (pancreas, liver) was excised and put into RNA*later*^TM^ stabilization solution (Invitrogen #2766377) and kept on ice or at -20 °C until extraction. This was necessary to ensure RNA integrity, especially from the pancreas, which has many RNAses which rapidly start degrading RNA at animal endpoint. RNA was extracted from mouse tissue using Trizol^TM^ (Invitrogen #15596018) according to the manufacturer’s protocol. In short, tissues were removed from RNA*later* and put into a chilled 2mL Eppendorf tube with 1mL of Trizol^TM^ and a stainless steel ball for tissue lysing (Grainger #4RJK6). Samples were lysed for 30 seconds at 25 Hz (Qiagen, TissueLyser) before 200 uL of chloroform was added and inverted several times. After 5 minutes, samples were centrifuged at max speed for 20 minutes at 4 °C and the clear supernatant (RNA fraction) was moved to a new tube. Isopropanol was used to precipitate the RNA and 75% ethanol was used to clean and dry the RNA pellet. Once the pellet was dry (roughly 10 minutes at room temp with lids opened), RNA was re-suspended in nuclease-free water (Promega #P119C). Samples were then stored at -80 °C until cDNA was generated.

Isolated RNA was thawed on ice and quantified using a NanoDrop 2000 (Thermo Fisher Scientific). cDNA was generated from 2 ug RNA of each sample following the protocol outlined with the High-Capacity cDNA Reverse Transcription Kit (Applied Biosystems #4368813). A Biometra T3 thermocycler was used for amplification. Following amplification, cDNA was diluted 1:20 in ddH_2_O. Once diluted, cDNA was used for qPCR analysis using *Power*SYBR® Green PCR Master Mix (Applied Biosystems #4367659) and following the manufacturer’s protocol. In short, 4 uL of diluted cDNA and 16 uL of PCR master mix were added to the well of a 384-well plate. Samples were loaded in triplicate and qPCR analysis was done using a Viia7 Real-time PCR System (Thermo Fisher Scientific).

QPCR data were analyzed by calculating the ΔΔCT as previously described^81^. In short, the CT value for each sample::gene pairing was normalized to housekeeping gene CTs which should not change across groups (TBP, 36b4, HPRT) and this value then normalized to the control animal group (ie. DBT^fl/fl^ group).

The forward and reverse primer pairs used for qPCR analysis were mixed for each gene of interest and their sequences are: mDBT (Fwd: AGA CTG ACC TGT GTT CGC TAT, Rev: GAG TGA CGT GGC TGA CTG TA); mDLD (Fwd: GAG CTG GAG TCG TGT GTA CC, Rev: CCT ATC ACT GTC ACG TCA GCC); mBCKDHe1a (Fwd: ATC TAC CGT GTC ATG GAC CG, Rev: ATG GTG TTG AGC AGC GTC AT); mBCKDHe1b (Fwd: AGC TAT TGC GGA AAT CCA GTT T, Rev: ACA GTT GAA AAG ATC ACC TGA GC); mBCKDK (Fwd: ACA TCA GCC ACC GAT ACA CAC, Rev: GAG GCG AAC TGA GGG CTT C); mBCAT1 (Fwd: CCC ATC GTA CCT CTT TCA CCC, Rev: GGG AGC GTG GGA ATA CGT G); mBCAT2 (Fwd: CTC ATC CTG CGC TTC CAG, Rev: TCA CAC CCG AAA CAT CCA ATC); mAMY2 (Fwd: CCT TCT GAC AGA GCC CTT GTG, Rev: GGA TGA TCC TCC AGC ACC AT); mLIP1 (Fwd: TTG TGG ACG CAA TTC ACA CA, Rev: TCT GGC TCA TTC CAA ATC CTA AG); mCTRB1 (Fwd: AAG AAC CCC AAG TTC AAC TCC TT, Rev: GGC AGG AGT GGC CAG CTT); mTRY3 (Fwd: TCT CTG AAC AGC GGC TAC CA, Rev: GTA GCA GTG AGC GGC TGA AAC); mINS1 (Fwd: TTC AGA CCT TGG CGT TGG A, Rev: AAA TGC TGG TGC AGC ACT GAT); mIRS1 (Fwd: TGA ACC TCA GTC CCA ACC ATA A, Rev: TCC GGC ACC CTT GAG TGT), mIRS2 (Fwd: GCC AGA AGG TGC CCG AGT, Rev: CCC CAG ATA CCT GAT CCA TGA); mTBP (Fwd: CCC TAT CAC TCC TGC CAC ACC AGC, Rev: GTG CAA TGG TCT TTA GGT CAA GTT TAC AGC C); m36b4 (Fwd: GGC TCC AAG CAG ATG CAG CAG, Rev: CCT GAT AGC CTT GCG CAT CAT GG); mHPRT (Fwd: GTT AAG CAG TAC AGC CCC AAA, Rev: AGG GCA TAT CCA ACA ACA AAC TT).

### RNA-sequencing and Analysis

Pancreata of 5 ½ week old male animals were processed for RNA as detailed above. All tissues were harvested at 1:30-2:00 PM to minimize variability. After RNA quality was confirmed using the NanoDrop 2000, RNA was sent to Novogene for standard non-directional library prep and bulk mRNA-sequencing using a NovaSeq PE150 (Ilumina) at a depth of 6Gb (20M PE reads) sequencing/sample. Fastq files were trimmed using Fastp^82^ and Salmon^83^ was used to count the trimmed data against the Gencode vM35 transcriptome, built on the GRCm39 genome. Several Bioconductor^84^ (v3.20) packages in R (v4.4.2) were used for subsequent steps. The transcriptome count data were annotated and summarized to the gene level with tximeta^85^ and further annotated with biomaRt^86^. PCA analysis and plots were generated with PCAtools^87^. Normalizations and statistical analyses were done with DESeq2^88^. QC and gene expression plots were performed with the DEGreport^89^ package.

Analysis for commonly downregulated and upregulated genes from bulk RNA-seq was done by filtering for genes that were significant hits in both comparisons: 1). DBT-pKO vs. DBT^fl/fl^ and 2). DBT-pKO vs. P48-Cre. Cutoffs to determine significance were (abs(log2ratio)) > 1.5. and a false discovery rate (FDR) adjusted p-value of ≤ 0.05. Data available for download on GEO: Accession #GSE299116.

### Western Blotting

Animals were euthanized in the afternoon and immediately the tissue of interest (pancreas, liver) was excised and put into 1mL of chilled RIPA buffer in a 2 mL Eppendorf tube. To this tube, a stainless steel ball for tissue lysing (Grainger #4RJK6) was added. Samples were lysed for 30 seconds at 20 Hz (Qiagen, TissueLyser) and the samples were then spun down at max speed for 20 minutes at 4 °C. This supernatant was saved and then quantified for lysate prep. To measure protein concentration in these tissue lysates, first 200 uL of tissue lysate was added to 800 uL of chilled RIPA buffer. RIPA buffer also contained 2X the normal concentration of both PhosSTOP (Roche 04906837001) and Complete PIC (Roche 82336300) to help neutralize many of the proteases and phosphatases that naturally start degrading sample after euthanasia. A BCA quantitation assay was then performed according to the manufacturer protocol (Thermo Scientific Pierce BCA Protein Assay #23228). An albumin standard curve (Thermo Scientific #23209) was run in parallel to allow for protein quantitation. Samples were then boiled with 4x Laemmli buffer for 10 minutes at 95 °C with 10% β-mercapto-ethanol (Sigma). These boiled samples were then frozen at -20 °C until they were run on a western blot.

For running western blots, 20 ug of boiled protein lysate was added per lane of a 10-, 12, or 16-lane 4-12% NuPAGE Bis/Tris pre-cast gels (Thermo Scientific Invitrogen). A BLUEstain protein standard ladder (Gold Biotechnology #P007-500) was run alongside samples for protein molecular weight estimation. Samples ran for 15 minutes at 100 V and then another 1 hour 15 minutes at 125 volts using NuPAGE MES SDS Running Buffer (Invitrogen NP0002) in a Mini Gel Tank (Invitrogen) powered by a PowerEase 300W (Life Technologies). Protein was then transferred onto 0.45uM nitrocellulose membranes (Bio-Rad #1620115) using a tris/glycine-based transfer buffer with 20% methanol running at room temperature for 1 hour at 20 volts. These membranes were then stained with Ponceau () to visualize protein loading and confirm even loading. These membranes were then blocked in 5% milk (in 1x TBST) for 1hour. 1X TBST was made with 10X TBST (Boston BioProducts #BM-300) and 0.1% Tween 20 (Bio-Rad #1706531). Samples were then washed with 1X TBST before they were incubated in primary antibody solution (3% BSA (Chem Cruz SC-2323) in 1XTBST) overnight rocking at 4°C. Primary antibody concentration used for all westerns was 1:1,000. The next morning, membranes were washed with 1X TBST on a rocker for 3 washes of 5 minutes each. LI-COR secondary antibodies were prepared in 3% BSA (in 1X TBST) at a concentration of 1:10,000 corresponding to the host in which the primary antibodies were raised in. Membranes were incubated in secondary solution for 1 hour at room temp on the rocker covered from light. Membranes were then washed three times with 1X TBST before being imaged on a Odyssey CLx (LI-COR).

The antibodies used for the western blots presented here were as follows with concentration used: HSP90 (Cell Signaling #4877; 1:2,500), HSP60 (Cell Signaling #12165; 1:2,500), DBT (Proteintech 12451-1-AP; 1:1,000), BCKDH-e1a (Cell Signaling #9011985; 1:1,000), BCKDH-e1b (Abcam ab201225; 1:1,000) DLD (Proteintech 16431-1-1AP; 1:1,000), s6K (Cell Signaling #9202; 1:1,000), p-s6k (Cell Signaling #9205; 1:1,000), 4E-BP1 (Cell Signaling #9644; 1:1,000), p-4E-BP1 (Cell Signaling #2855; 1:1,000), s6 (Cell Signaling #2317; 1:1,000), p-s6 (Cell Signaling #2211; 1:1,000), AMPK-a (Cell Signaling #25325; 1:1,00), p-AMPK-a (Cell Signaling #2535; 1:1,000), BCAT2 (Invitrogen PAS-21549; 1:1,000), BCKDK (Sigma HPA-017995; 1:1,000), p-BCKDH-e1a (Thermo Scientific A304-672A; 1:1,000), PDH-e1a (Cell Signaling 3205T; 1:1,000), p-PDH-e1a (Cell Signaling #31866; 1:1,000), OGDH (Proteintech 15212-1-AP; 1:1,000), PPM1K (Proteintech 14573-1-AP; 1:1,000). Secondary antibodies were: IRDye 680 Goat anti-Ms (LI-COR 926-68070, 1:10,000) and IRDye Goat anti-Rb (LI-COR 926-32210, 1:10,000).

### Pancreatic Tumor Initiation and Scoring

Animals from tumor-prone mouse models (KC, KC;DBT-pKO, KC;BCAT2-pKO, KC;DBT,BCAT2-pKO, and KC;BCKDK-pKO) were enrolled into tumor-initiating assays at birth. Animals were weighed weekly starting at 4 weeks old until endpoint. For cerulein-initiating tumor assays, animals were injected at 6 ½ weeks old. Cerulein (Sigma C9026) was injected at 50 ug/kg intra-peritoneally 4 times per day (every 2 hours) for 2 consecutive days (Day 0 and Day 1). At Day 9, mice were euthanized and pancreata are harvested for histology collection and prepped as described above. For non-cerulein-based tumor assays, animals were euthanized at either 8 or 12 weeks old and their pancreata were harvested for histology collection. HCE slides of pancreata from tumor-bearing animals were all imaged on a Keyence as previously mentioned and normal acinar area was calculated.

8 HCE slides of animals with healthy acinar area closest to the mean for each group were submitted to the Comparative Pathology Core in the School of Veterinary Medicine at the University of Pennsylvania for analysis. Blinded histopathological evaluation and classification of the proliferative and non-proliferative lesions observed in the KC mouse models of multistep pancreatic carcinogenesis were performed according to consensus criteria established at the Penn Workshop and the subsequent MMHCC consensus report^90^. The histopathological examination has confirmed the typical multicentric development of mouse pancreatic intraepithelial neoplasia (PanIN) in all of the examined samples with varying degrees of severity in terms of number, extension, and grade of the lesions. A multiparametric scoring system for ADM and inflammation was used to assess changes in pancreatic carcinogenesis. ADM scoring consisted of 3 categories each with a possible score of 0-3 with the aggregate score reported here. Inflammation scoring was scored from 0-3 with that score reported here. A breakdown of the ADM scoring criteria used for assessment is reported in Extended Data Fig. 3c and follows here:

1. ADM extension and distribution (0 = no ADM observed; 1 = ADM involving up to 10% of the pancreatic section; 2= ADM involving from 11% up to 30% of the pancreatic section; 3 = ADM involving more than 30% of the pancreatic section)
2. Overall atypical features associated with ADM (0 = no ADM observed; 1 = ADM mainly characterized by nonatypical features; 2 = ADM characterized by both atypical and nonatypical features; 3 = ADM mainly characterized by atypical features)
3. Overall level of fibrosis associated with ADM (0 = no ADM observed; 1 = mild, fibrosis mainly confined to the periphery of the affected lobule/group of acini; 2 = moderate, fibrosis that from the periphery of the affected lobule/group of acini separates the individual acinar profiles; 3 = severe, characterized by a dense fibrotic nodule with few scattered ADM profiles).

Inflammation Multiparametric Scoring Criteria is presented in Extended Data Fig. 3d and was as follows: (0 = no inflammation observed; 1 = minimal inflammatory cell infiltrates confined to the fibrotic interstitium surrounding ADM; 2 = moderate inflammatory cell infiltrates accompanying ADM and extending into the fibrotic/edematous interlobular interstitium; 3 = severe inflammatory cell infiltrates accompanying ADM, extending the interlobular interstitium, and involving the fibrotic/edematous pancreatic capsule).

### Intratracheal Cre delivery and Lung Tumor Formation

Animals with floxed alleles (LSL-Kras^G12D^, LSL-Kras^G12D^;DBT^fl/fl^ and LSL-Kras^G12D^;BCKDK^fl/fl^) and no Cre were used for lung tumor formation assays. At 6-10 weeks old, tumors were transduced via endotracheal inhalation of a Cre-recombinase expressing adenovirus (Ad:CMV-Cre) as previously described^91^. Animals were transduced with 2.5x 10^5^ plaque forming units (PFU) per mouse in 50 uL volume injections after they were placed under general anesthesia (isoflurane). Mice were then placed on a heating pad and monitored until they were fully recovered. Animals were monitored for 20 weeks until endpoint.

At endpoint, lungs were inflated using a 10% neutral-buffered formalin solution at time of harvest and were fixed at room temperature overnight. The next day, lungs were dehydrated in a graded alcohol series similar as to that described above for pancreata. Tissues were ultimately paraffin-embedded by the Penn Molecular Pathology and Imaging Core and HCE sections were produced. Immunohistochemistry was performed on paraffin-embedded sections using a Leica BOND RXm Automatic Slide Stainer with a Leica BOND Polymer Refine Detection kit (Leica Biosystems, BOND Polymer Refine Detection Kit, DS9800). Following deparaffinization and citrate heat induced antigen retrieval, sections were incubated with the following antibody for 30 minutes at room temperature: Ki-67 (1:800, Novus, #89717). Sections were then incubated with a poly-HRP anti-rabbit IgG polymer (Leica Biosystems, BOND Polymer Refine Detection Kit, DS9800) for 8 minutes at room temperature. Signal was developed with DAB chromogenic substrate (Leica Biosystems, BOND Polymer Refine Detection kit, DS9800) followed by counterstaining with hematoxylin (Leica Biosystems, BOND Polymer Refine Detection Kit, DS9800). Whole-slide scans were generated using a Leica DMI6000B microscope at 5x magnification. Via these scans, staining was quantified on ImageJ using Mean Gray Value as previously described^92^. All mean gray values were then divided by 255 to convert the value to percent positive area.

### BT2 Chow Formulation

3,6-Dichlorobenzo(b)thiophene-2-carboyxlic acid (BT2, Chem Impex #25643) was sent to Research Diets, Inc. where it was incorporated into the AIN-76A diet (Research Diets D10001) at 250 mg/kg as previously described^45^. BT2 and control AIN-76A diets were provided to the animals *ad libitum* and were changed out every 3 days.

### Data Analysis

All data were analyzed and processed in Microsoft Excel and graphed using GraphPad Prism 10. Statistical analyses were done using GraphPad Prism and specific statistical tests are included in figure legends. Natural abundance correction for isotope tracing experiments was performed using FluxFix and unlabeled controls were used to normalize labeling.

## Acknowledgements

We would like to thank the University of Pennsylvania’s Center for Molecular Studies in Digestive and Liver Diseases (P30DK050306) and the Molecular Pathology and Imaging Core (RRID:SCR_022420) for performing tissue mounting and HCE staining. We would also like to thank the Comparative Pathology Core in the School of Veterinary Medicine at the University of Pennsylvania for performing histological evaluation of mouse pancreata. Further we would like to thank all Arany and Wellen lab members for critical feedback and discussion, with particular recognition to Michael Neinast, Alessandro Carrer, Megan Blair, Wencao Zhao, and Julianna Supplee for discussion; and Jessie Axsom, Nicholas Forelli, and Jae Woo Jung for LC-MS troubleshooting. Lastly, we would like to acknowledge the use of BioRender for figure generation.

---

**Extended Data Figure 1.**
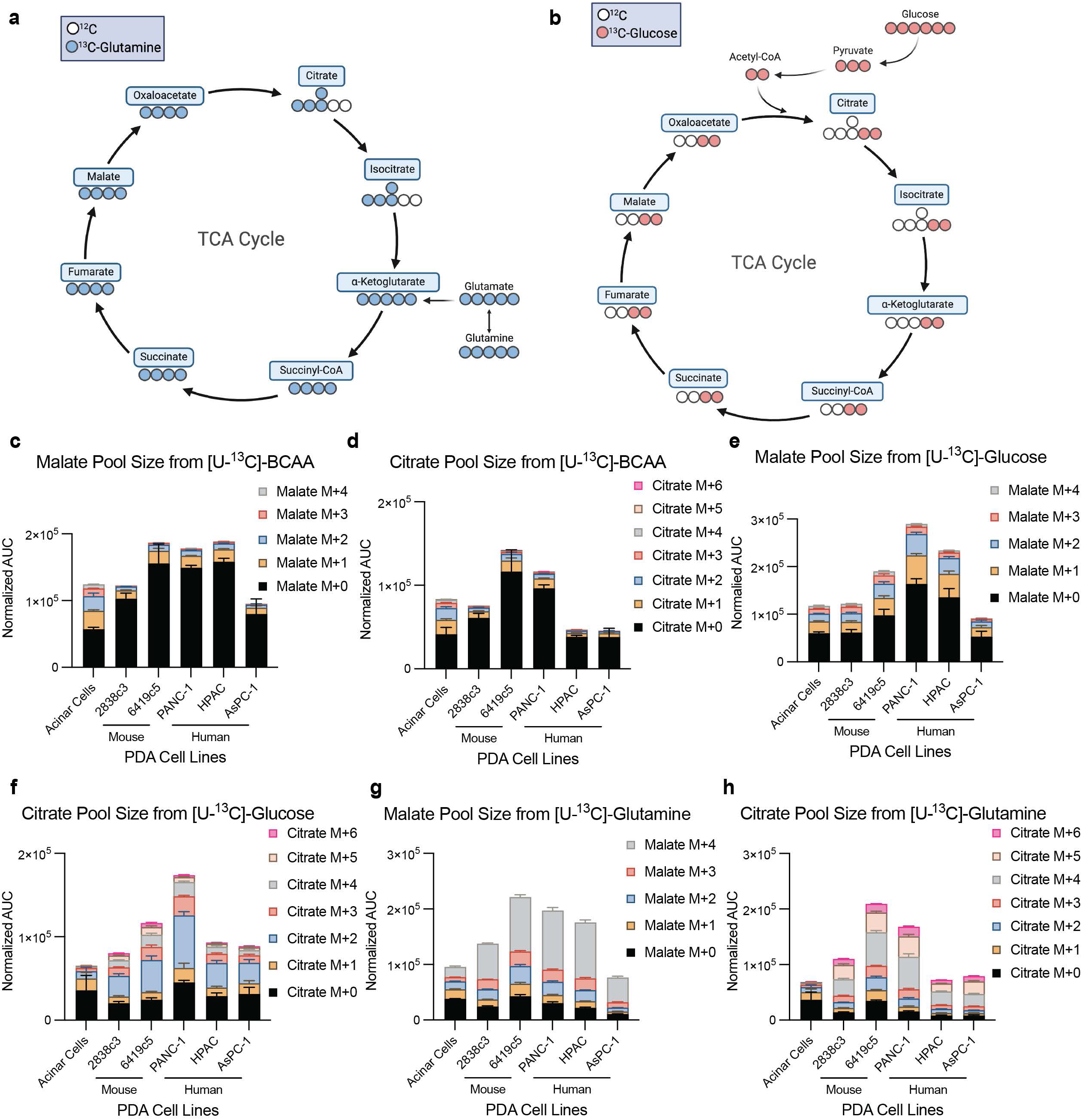
Contribution of nutrients to the TCA cycle in acinar and PDA cells. **a**, Tracing schematic for U-^13^C-glutamine incorporation into TCA cycle intermediates via α-ketoglutarate. **b**, Tracing schematic for U-^13^C-glucose incorporation into TCA cycle intermediates via acetyl-CoA. **c-h**, Malate and citrate pool size with individual isotopologues matching MPE data in Fig. 1d-i. Abundance is normalized to cell number (n=3 biological replicates per cell line). Data are presented as mean ± s.e.m.

**Extended Data Figure 2.**
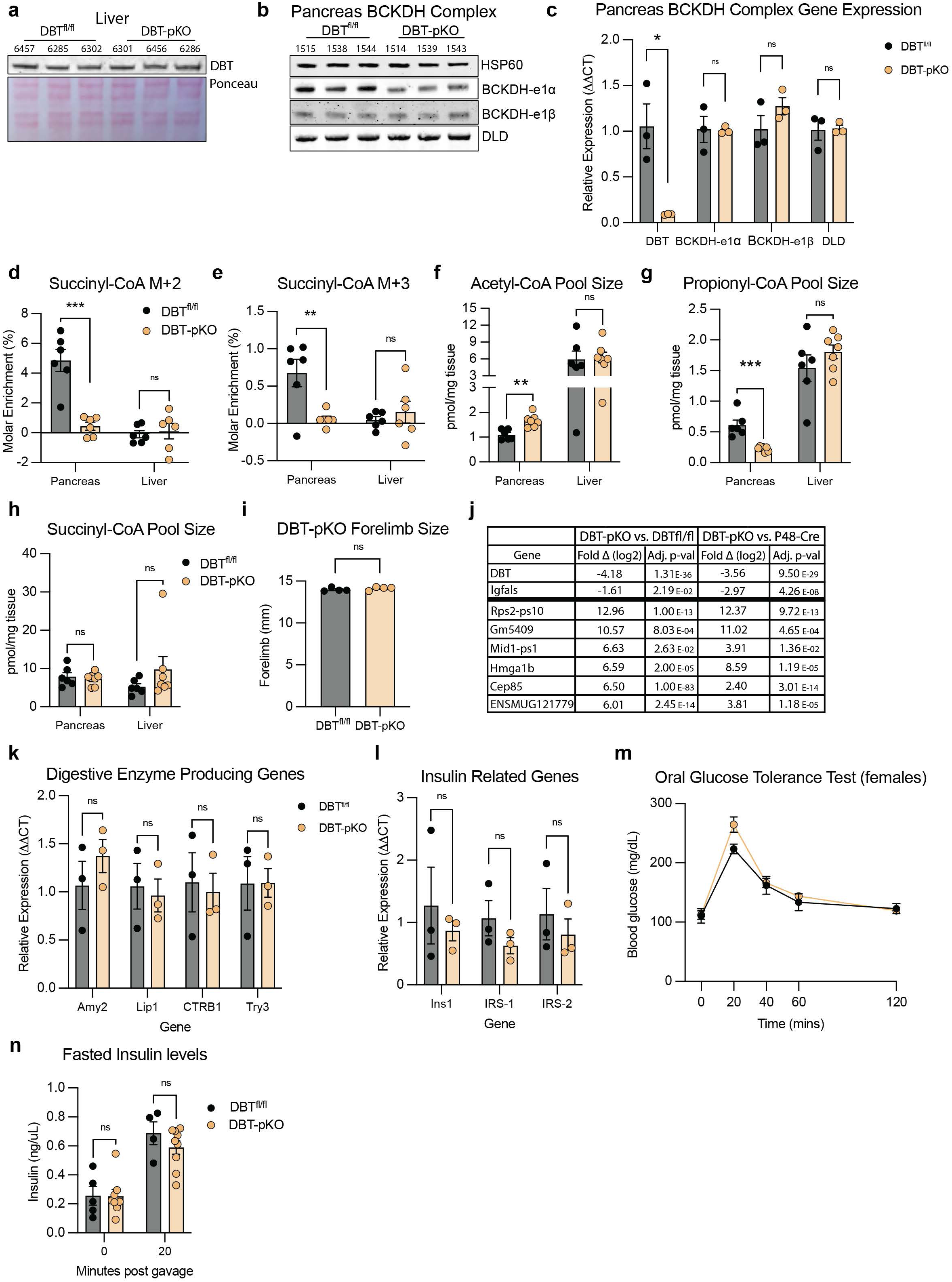
Characterization of DBT-pKO animals. **a**, Western for DBT protein in liver of 8-10-week-old animals from Fig. 2a (n=3 DBT^fl/fl^ and n=3 DBT-pKO; Ponceau is the loading control). **b**, Western for BCKDH complex subunit proteins (p-BCKDH-e1α, BCKDH-e1α, DLD) in pancreata of 8-10-week-old animals (n=3 DBT^fl/fl^ and n=3 DBT-pKO; HSP60 is the loading control). **c**, qPCR of pancreata extracted from 8-10-week-old animals showing BCKDH complex gene expression (DBT, BCKDH-e1α, BCKDH-e1β, and DLD) (n=3 DBT^fl/fl^ and n=3 DBT-pKO). Data presented as ΔΔCT and normalized to DBT^fl/fl^ for relative expression. **d,e**, Succinyl-CoA M+2 and succinyl-CoA M+3 MPE measured in animal pancreata and liver 20 mins after bolus gavage of [U-^13^C]-BCAAs as introduced in 2b (n=6 DBT^fl/fl^ and n=6 DBT-pKO). **f-h**, Pool size for acetyl-CoA, propionyl-CoA, and succinyl-CoA with stacked isotopologues matching MPE from Fig. 2c,d and Extended Data Fig. 2d,e (n=6 DBT^fl/fl^ and n=6 DBT-pKO). **i**, Forelimb size measured at endpoint in 12-week-old animals with calipers (n=4 DBT^fl/fl^ and n=4 DBT-pKO). **j**, Table of significant genes from Fig. 2g,h with corresponding log2 fold change and adjusted p-values listed for both group comparisons. **k,l** qPCR of pancreata extracted from 8-10-week-old animals showing digestive enzyme producing genes and insulin related genes (n=3 DBT^fl/fl^ and n=3 DBT-pKO). **m**, Oral glucose tolerance tests conducted on female littermates of animals presented in Fig. 2m (n=8 DBT^fl/fl^ and n=13 DBT-pKO). **n**, Insulin levels measured at 20 mins post oral gavage from animals in panel **m** (n=5 DBT^fl/fl^ and n=9 DBT-pKO). Data are presented as mean± s.e.m. When two groups are compared, a two-tailed Student’s t-test was used with significance defined as \**P* < 0.05, \*\**P* < 0.01, and \*\*\**P* < 0.001. **c**, DBT *P* = 0.0172. **d**, Succinyl-CoA M+2 *P*=0.0002. **e**, Succinyl-CoA M+3 *P*=0.0084. **f**, Acetyl-CoA pool size pancreas *P*=0.0012. **g**, Propionyl-CoA pool size pancreas *P*=0.0003.

**Extended Data Figure 3.**
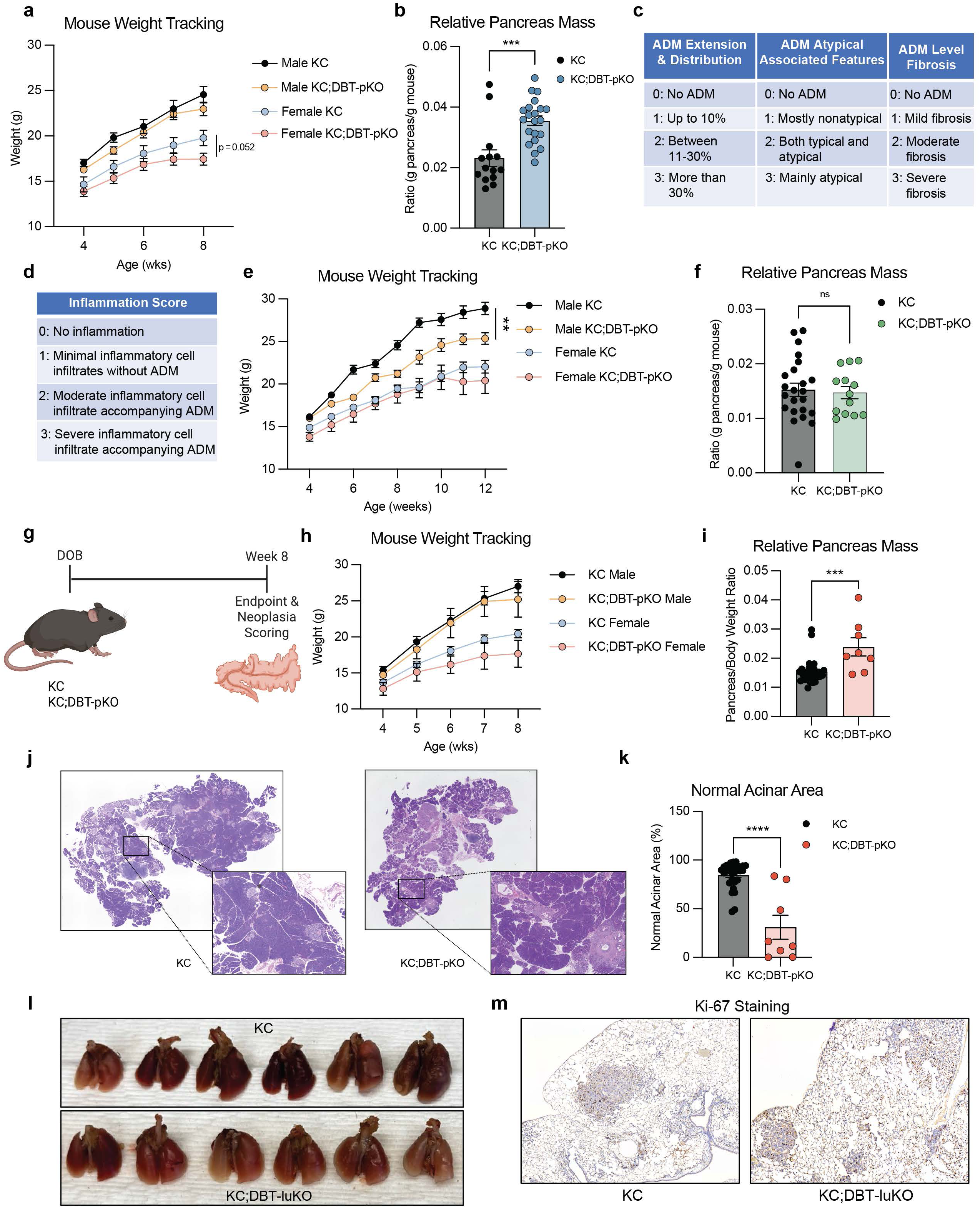
Further characterization of KC;DBT-pKO animals and lung DBT KO tumor initiation model. **a**, Mouse weight recorded weekly and shown separated by sex (n=6 male KC, n=10 male KC;DBT-pKO, n=6 female KC, and n=11 female KC;DBT-pKO). Statistical value shown for 8-week measurement only. **b**, Relative pancreas mass measured by normalizing pancreas weight at endpoint to total body weight corresponding to Fig. 3c (n=14 KC and n=21 KC;DBT-pKO). **c,d**, Multi-parametric scoring criteria tables for ADM score shown in Fig. 3e and inflammation score shown in Fig. 3f. **e**, Mouse weight recorded weekly for 12 weeks and shown separated by sex (n= 12 male KC, n=8 male KC;DBT-pKO, n=11 female KC, and n=5 female KC;DBT-pKO). **f**, Relative pancreas mass measured by normalizing pancreas weight at 12-week endpoint to total body weight corresponding to Fig. 3i (n=23 KC and n=13 KC;DBT-pKO). **g**, Schematic and timeline of KC and KC;DBT-pKO animals euthanized at 8 weeks old without cerulein administration. **h**, Mouse weight recorded weekly and shown separated by sex (n=10 male KC, n=15 female KC, n=4 male KC;DBT-pKO, and n=4 female KC;DBT-pKO). **i**, Relative pancreas mass measured by normalizing pancreas weight at endpoint to total body weight corresponding to panel **h** (n=30 KC and n=8 KC;DBT-pKO). **j**, Representative HCE images of KC and KC;DBT-pKO pancreata collected at an 8-week timepoint. **k**, Normal acinar area was calculated by the total area of healthy acinar tissue relative to total pancreas area for each animal (n=30 KC and n=8 KC;DBT-pKO). **l**, Representative lung images from mice detailed in Fig. 3g at time of harvest after 20 weeks post-Cre administration (n=6 KC and n=6 KC;DBT-luKO). **m**, Representative Ki-67 staining image from lung lesions quantified in Fig. 3m. Data are presented as mean± s.e.m. Representative images depict animals closest to the mean of each group. When two groups are compared, a two-tailed Student’s t-test was used, with significance defined as \*\*\**P*<0.001 and \*\*\*\**P*<0.0001. **a**, 8-week female *P* =0.052. **b**, Relative pancreas mass *P*=0.0002. **e**, 12-week male weight *P*=0.0033. **i**, Relative pancreas mass *P*=0.0003. **k**, Normal acinar area *P*<0.0001.

**Extended Data Figure 4.**
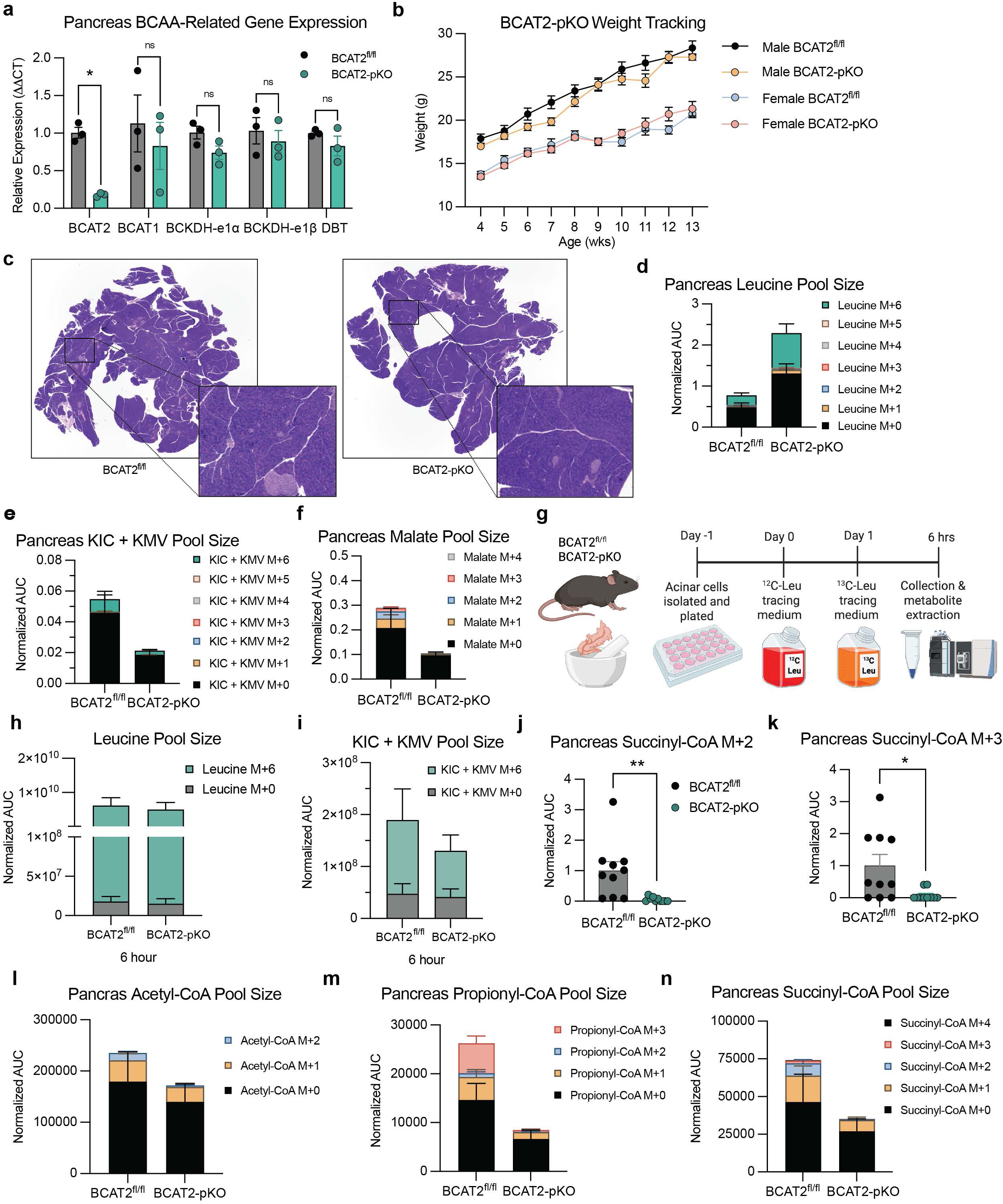
Further characterization of BCAT2-pKO animals. **a**, qPCR of pancreata from 8-week-old animals for BCAA-related gene expression (n=3 BCAT2^fl/fl^ and n=3 BCAT2-pKO). **b**, Animal weight recorded weekly and shown separated by sex (n=8 male BCAT2^fl/fl^, n=10 male BCAT2-pKO, n=8 female BCAT2f^l/fl^, and n=8 female BCAT2-pKO). **c**, Representative H+E staining image. **d-f**, Pool size for leucine, KIC+ KMV, and malate in pancreata of animals after [U-^13^C]-BCAA gavage with stacked isotopologues matching MPE data from Fig. 4e-g (n=10 BCAT2^fl/fl^ and n=10 BCAT2-pKO). **g**, Schematic and diagram for *ex vivo* acinar cell tracing of freshly isolated acinar cells from pancreata of animals provided [U-^13^C]-leucine for 6 hours prior to harvest for LC-MS analysis. **h,i**, M+0 and M+6 leucine and KIC isotopologues were measured after 6 hours of tracing (n=6 BCAT2^fl/fl^ and n=6 BCAT2-pKO). **j,k**, Succinyl-CoA M+2 and succinyl-CoA M+3 labeling, respectively, in bolus BCAA-provided BCAT2-pKO animals from Fig. 4d. **l-n**, Pool size for acetyl-CoA, propionyl-CoA, and succinyl-CoA from gavage matching MPE from Fig. 4h,i and Extended Data Fig. 4j,k (n=10 BCAT2^fl/fl^ and n=10 BCAT2-pKO). Data are presented as mean± s.e.m. When two groups are compared, a two-tailed Student’s t-test was used, with significance defined as \**P*<0.05 and \*\**P*<0.01. **g**, Succinyl-CoA M+2 *P* =0.0049. **h**, Succinyl CoA M+3 *P*=0.017.

**Extended Data Figure 5.**
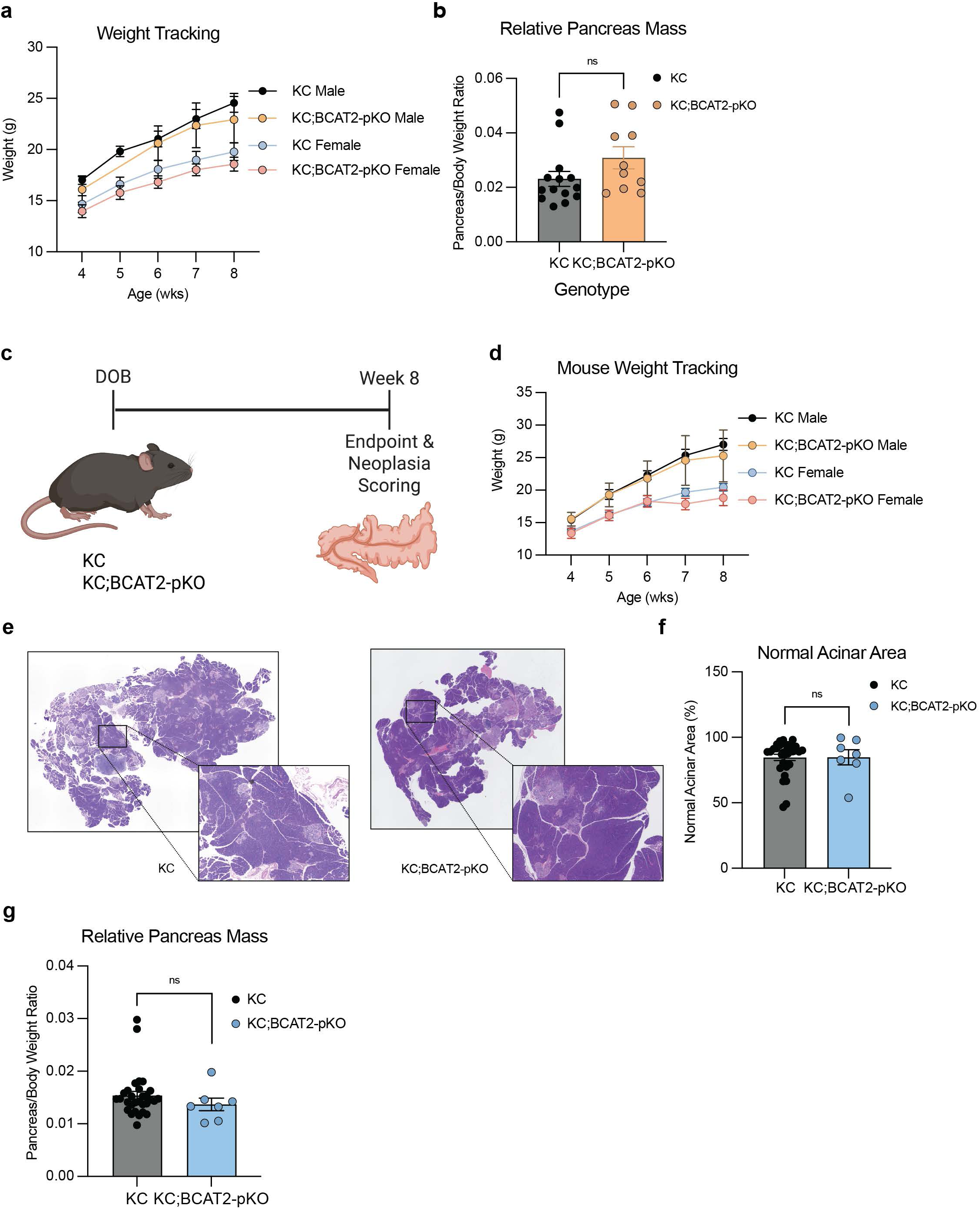
Further characterization of KC;BCAT2-pKO tumor initiation assays. **a**, Animal weight recorded weekly of animals euthanized at 8 weeks old with cerulein administration at 6 ½ weeks old. Data is shown separated by sex (n=8 male KC, n=3 male KC;BCAT2-pKO, n=6 female KC, and n=7 female KC;BCAT2-pKO). **b**, Relative pancreas mass measured by normalizing pancreas weight at endpoint to total body weight corresponding to Fig. 4m (n=14 KC and n=10 KC;BCAT2-pKO). **c**, Schematic and timeline for collection and analysis of 8-week-old KC and KC;BCAT2-pKO animals without cerulein administration. **d**, Animal weight recorded weekly of animals euthanized at 8 weeks old (n=10 male KC, n=15 female KC, n=3 male KC;BCAT2-pKO, and n=4 female KC;BCAT2-pKO. **e**, Representative H+E staining images show mouse pancreata at endpoint. **f**, Normal acinar area of animals harvested at 8 weeks with no inflammation insult (n=30 KC and n=7 KC;BCAT2-pKO). **d**, Representative H+E staining images show mouse pancreata at endpoint with red box around tumor. **e**, Animal weight recorded weekly of animals euthanized at 8 weeks old. Data is shown separated by sex (n=10 male KC, n=3 male KC;BCAT2-pKO, n=15 female KC, and n=4 female KC;BCAT2-pKO). **f**, Normal acinar area was calculated by the total area of healthy acinar tissue relative to total pancreas area for each animal (n=30 KC and n=7 KC;BCAT2-pKO). KC historical data presented in Extended Data Fig. 3g re-graphed for comparison purposes. **g**, Relative pancreas mass measured by normalizing pancreas weight at endpoint to total body weight corresponding to panel **d** (n=30 KC and n=7 KC;BCAT2-pKO). Data are presented as mean± s.e.m. Representative images depict animals closest to the mean of each group. When two groups are compared, a two-tailed Student’s t-test was used.

**Extended Data Figure 6.**
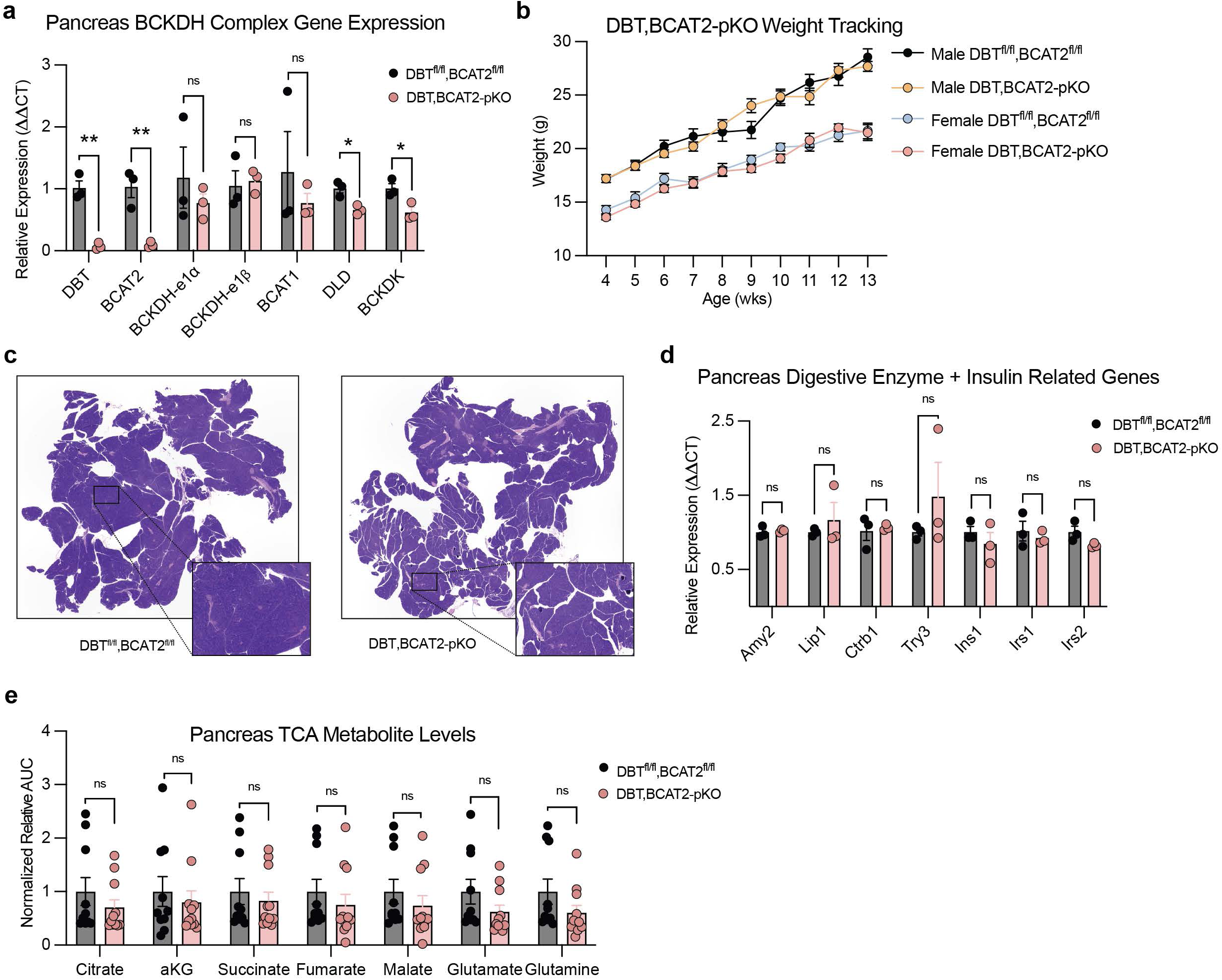
Further characterization of DBT,BCAT2-pKO animals. **a**, qPCR of pancreata from 8-week-old animals for BCAA-related gene expression (n=3 DBT^fl/fl^,BCAT2^fl/fl^ and n=3 DBT,BCAT2-pKO). **b**, Animal weight recorded weekly of animals euthanized at 12 weeks old. Data are shown separated by sex (n=9 male DBT^fl/fl^,BCAT2^fl/fl^, n=10 male DBT,BCAT2-pKO, n=8 female DBT^fl/fl^,BCAT2^fl/fl^, and n=9 female DBT,BCAT2-pKO). **c**, Representative H+E staining images show mouse pancreata at endpoint. **d**, qPCR of pancreata from 8-week-old animals for digestive enzyme and insulin related genes (n=3 DBT^fl/fl^,BCAT2^fl/fl^ and n=3 DBT,BCAT2-pKO). **e**, LC-MS relative measurements of pancreatic TCA metabolites (n=10 DBT^fl/fl^,BCAT2^fl/fl^ and n=11 DBT,BCAT2-pKO). Data are presented as mean± s.e.m. Representative images depict animals closest to the mean of each group. When two groups are compared, a two-tailed Student’s t-test was used for each individual gene, with significance defined as \**P*< 0.05 and \*\**P*< 0.01. **c**, DBT *P*=0.0015; BCAT2 *P*=0.0061; DLD *P*=0.0139; BCKDK *P*=0.0274.

**Extended Data Figure 7.**
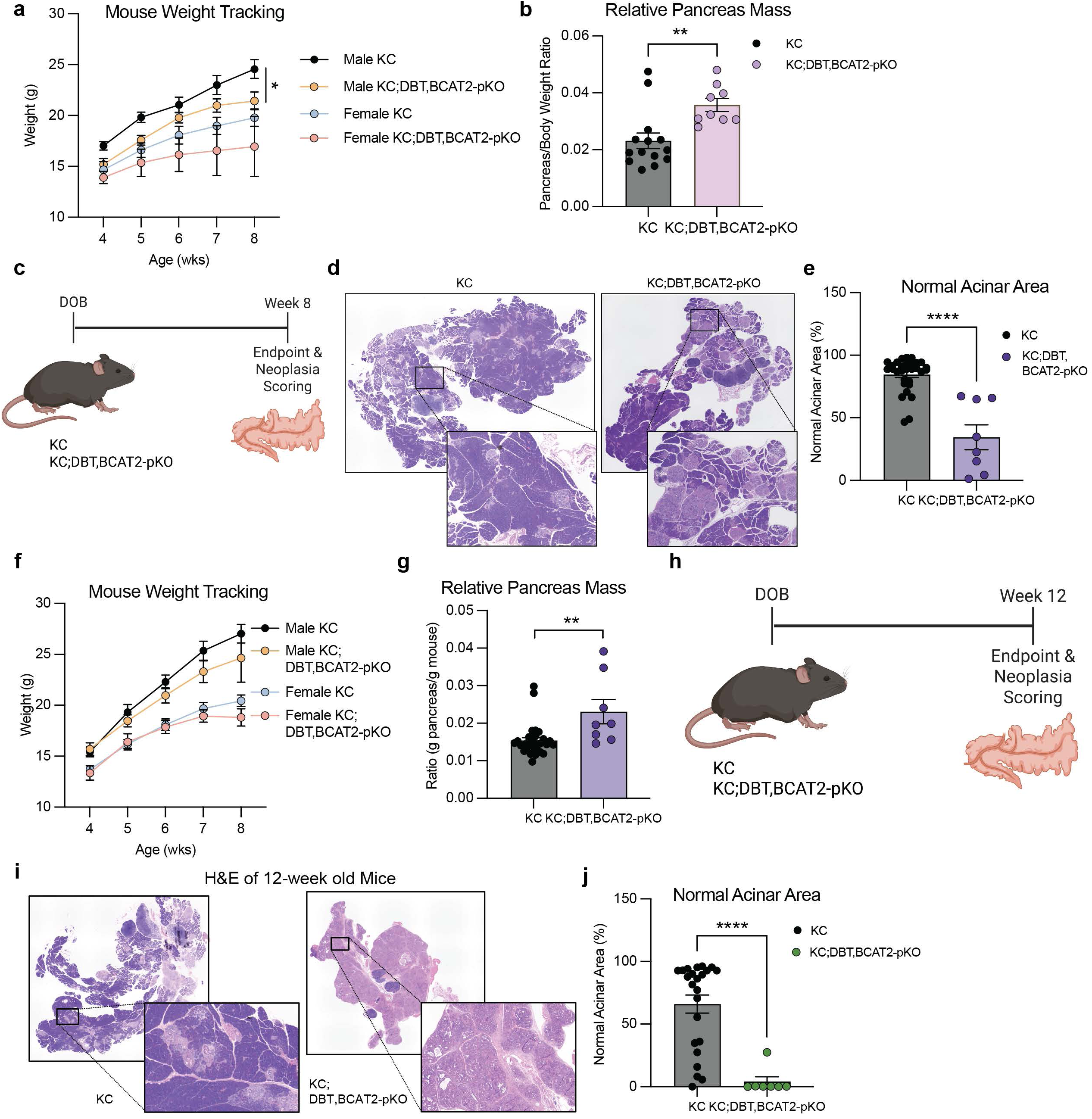
Further characterization of KC;DBT,BCAT2-pKO tumor initiation assays. **a**, Animal weight recorded weekly of animals euthanized at 8 weeks with cerulein administration at 6 ½ weeks old for animals shown in Fig 4q (n=8 male KC, n=7 male KC;DBT,BCAT2-pKO, n=6 female KC, and n=2 female KC;DBT,BCAT2-pKO). **b**, Relative pancreas mass measured by normalizing pancreas weight at endpoint to total body weight corresponding to Fig. 4s (n=14 KC and n=9 KC;DBT,BCAT2-pKO). **c**, Schematic and timeline for KC;DBT,BCAT2-pKO 8-week tumor initiation assays. **d**, Representative H+E images of pancreata from KC and KC;BCAT2-pKO animals at endpoint. **e,** Normal acinar area of animals harvested at 8 weeks with no inflammation insult (n=30 KC and n=8 KC;DBT,BCAT2-pKO). KC historical data presented in Extended Data Fig. 3g re-graphed for comparison purposes. **f**, Animal weight recorded weekly of animals euthanized at 8 weeks for animals shown in panel **c** (n=10 male KC, n=3 male KC;DBT,BCAT2-pKO, n=15 female KC, and n=5 female KC;DBT,BCAT2-pKO). **g**, Relative pancreas mass measured by normalizing pancreas weight at endpoint to total body weight corresponding to panel **f** (n=30 KC and n=8 KC;DBT,BCAT2-pKO). **h**, Schematic and timeline for KC;DBT,BCAT2-pKO 12-week tumor initiation assays. **i**, Representative HCE images of pancreata at endpoint. **j**, Normal acinar area of animals harvested at 12 weeks with no inflammation insult (n=23 KC and n=7 KC;DBT,BCAT2-pKO). KC historical data presented in Fig. 3g re-graphed for comparison purposes. Data are presented as mean± s.e.m. Representative images depict animals closest to the mean of each group. When two groups are compared, a two-tailed Student’s t-test was used, with significance defined as \**P*< 0.05, \*\**P*< 0.01 and \*\*\*\**P*< 0.0001. **a**, 8-week male endpoint *P*=0.0296. **b**, Relative pancreas mass *P*=0.0039. **e**, Normal acinar area *P*<0.0001. **g**, Relative pancreas mass *P*=0.0012. **j**, Normal acinar area *P*<0.0001.

**Extended Data Figure 8.**
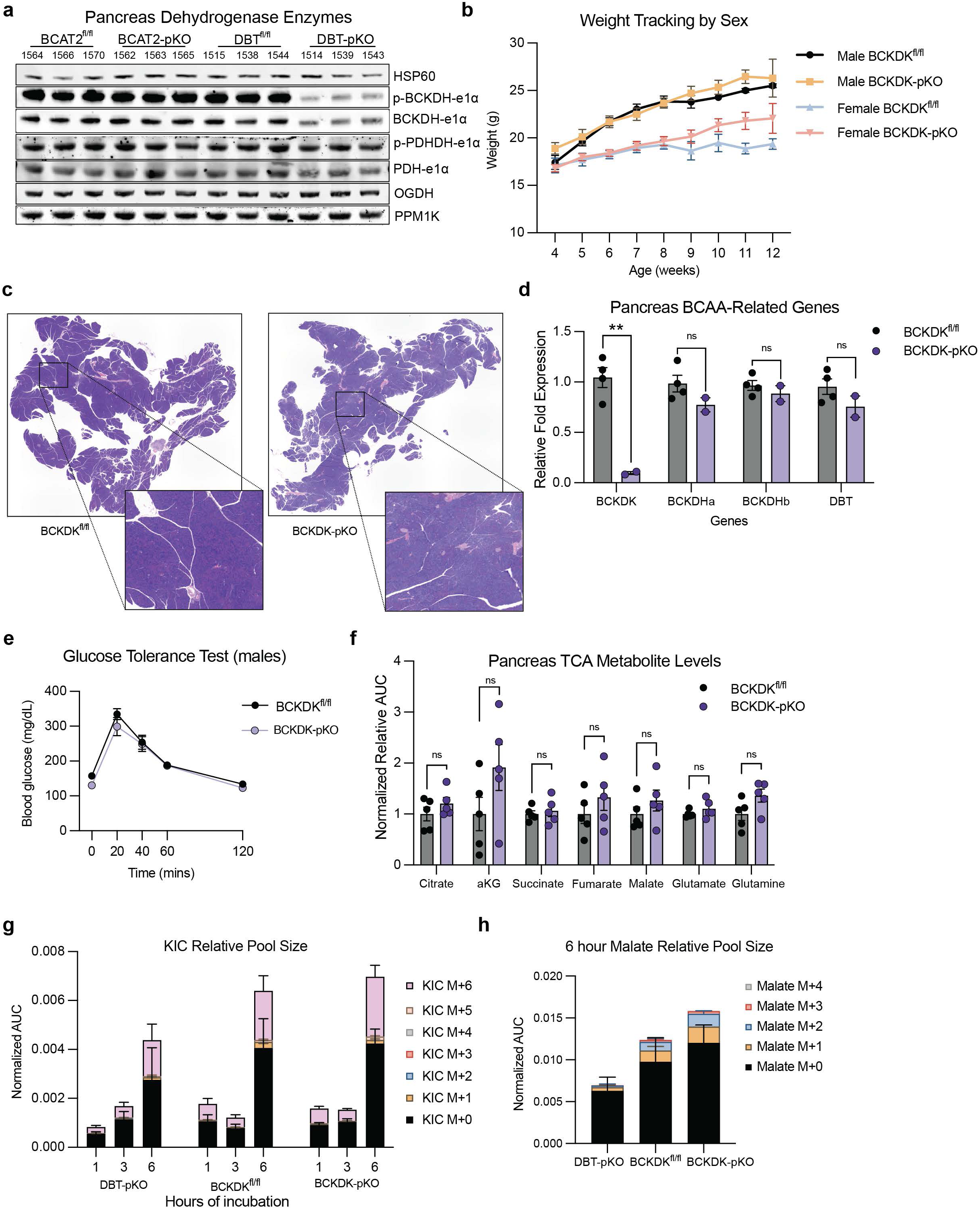
Further characterization of BCKDK-pKO animals. **a**, Western blots in pancreata for various dehydrogenase enzymes from indicated genotypes of 8-week-old animals (n=3 per genotype; HSP60 is the loading control). **b**, Animal weight recorded weekly of animals euthanized at 12 weeks old. Data are shown separated by sex (n=19 male BCKDK^fl/fl^, n=19 male BCKDK-pKO, n=11 female BCKDK^fl/fl^, n=18 BCKDK-pKO). **c**, Representative H+E images. **d**, qPCR of pancreata extracted from 8-week-old animals for BCAA-related genes (n= 4 BCKDK^fl/fl^ and n=2 BCKDK-pKO). **e**, Oral glucose tolerance tests conducted on animals 8-week-old male animals (n=8 BCKDK^fl/fl^ and n=8 BCKDK-pKO). **f**, LC-MS relative measurements of pancreatic TCA metabolites (n=5 DBT^fl/fl^,BCAT2^fl/fl^ and n=5 DBT,BCAT2-pKO). **g**, Pool size for KIC shown after 1, 3, and 6 hours of incubation with [U-^13^C]-leucine with stacked isotopologues matching labelling from Fig. 5g (n=6 DBT-pKO, n=4 BCKDK^fl/fl^, and n=3 BCKDK-pKO). **h**, Pool size for malate shown after 6 hours of labelling with stacked isotopologues matching labelling from Fig. 5h (n=6 DBT-pKO, n=4 BCKDK^fl/fl^, and n=3 BCKDK-pKO). Data are presented as mean± s.e.m. Representative images depict animals closest to the mean of each group. When two groups are compared, a two-tailed Student’s t-test was used, with significance defined as \*\**P*< 0.01. **g**, BCKDK *P* =0.0030.

**Extended Data Figure 9.**
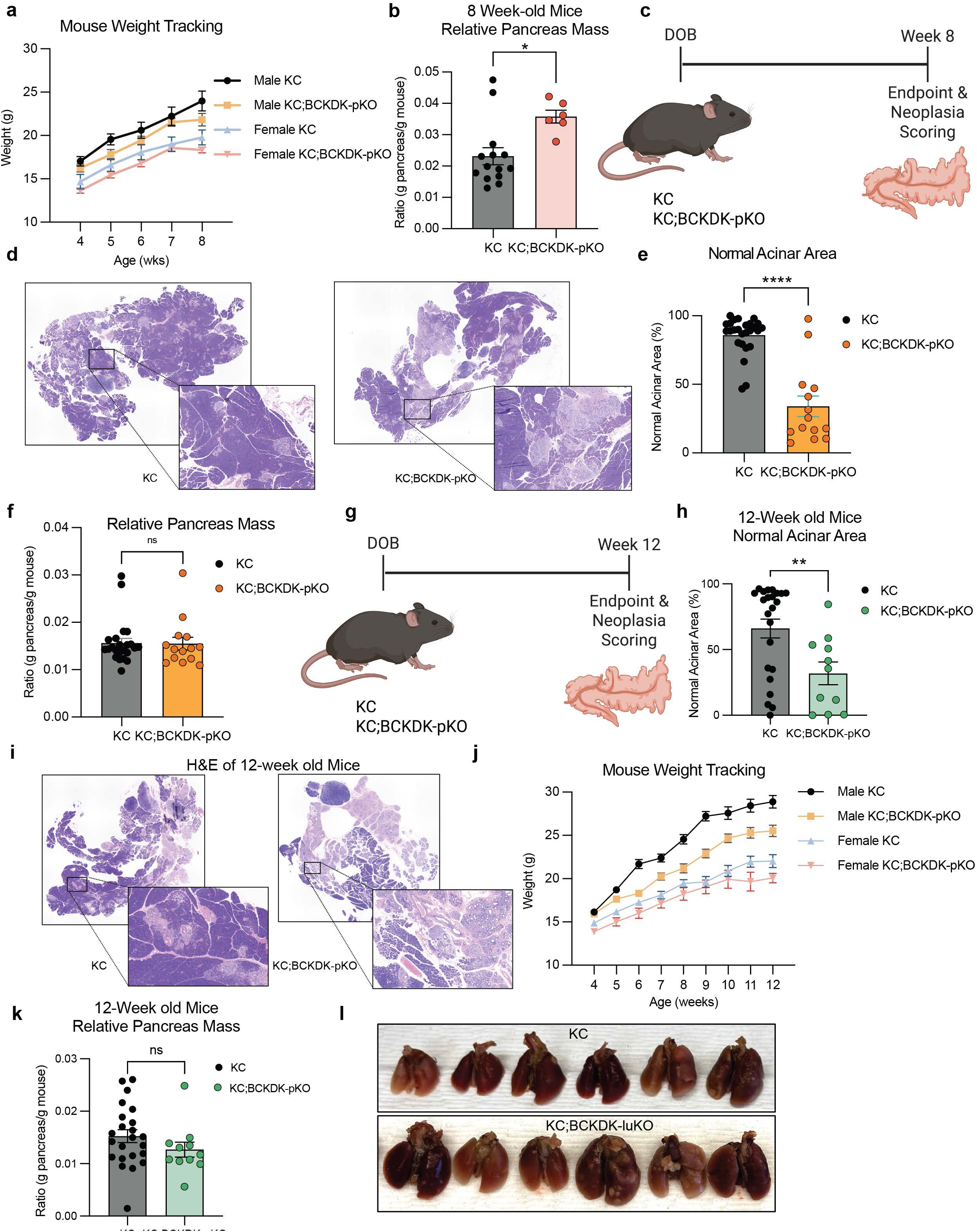
KC;BCKDK-pKO animals exhibit increased PanIN formation. **a**, Animal weight recorded weekly of animals euthanized at 8 weeks old with cerulein administration at 6 ½ weeks. Data is shown separated by sex (n=6 male BCKDK^fl/fl^, n=3 male BCKDK-pKO, n=6 female BCKDK^fl/fl^, n=3 BCKDK-pKO). **b**, Relative pancreas mass measured by normalizing pancreas weight at endpoint to total body weight corresponding to Fig. 5k (n=14 KC and n=6 KC;BCKDK-pKO). **c**, Schematic and timeline for 8-week tumor initiation assay without cerulein administration. **d**, Representative H+E staining images show mouse pancreata at endpoint. **e**, Normal acinar area was plotted relative to total pancreas area for animals euthanized at 8 weeks old (n=26 KC and n=14 KC;BCAT2-pKO). **f**, Relative pancreas mass measured by normalizing pancreas weight at endpoint to total body weight corresponding to Ext. Fig. 9e (n=26 KC and n=14 KC;BCAT2-pKO). **g**, Schematic and timeline for 12-week tumor initiation assay without cerulein administration. **h**, Normal acinar area was plotted relative to total pancreas area for animals euthanized at 8 weeks old (n=23 KC and n=11 KC;BCAT2-pKO). **i**, Representative H+E staining images show mouse pancreata at endpoint. **j**, Animal weight recorded weekly of animals euthanized at 12 weeks old. Data is shown separated by sex (n=12 male BCKDK^fl/fl^, n=12 male BCKDK-pKO, n=11 female BCKDK^fl/fl^, n=8 BCKDK-pKO). **k**, Relative pancreas mass measured by normalizing pancreas weight at endpoint to total body weight corresponding to panel **h** (n=14 KC and n=9 KC;DBT,BCAT2-pKO). **l**, Representative images of lungs from mice detailed in Fig. 5l at time of harvest (n=6 KC and n=6 KC;BCKDK-luKO. Data are presented as mean± s.e.m. Representative images depict animals closest to the mean of each group. When two groups are compared, a two-tailed Student’s t-test was used, with significance defined as \**P* <0.05 and \*\*\*\**P*<0.001. **b**, Relative pancreas mass *P*=0.0108. **e**, Normal acinar area *P* <0.0001. **h**, Normal acinar area *P*=0.0075.

**Extended Data Figure 10.**
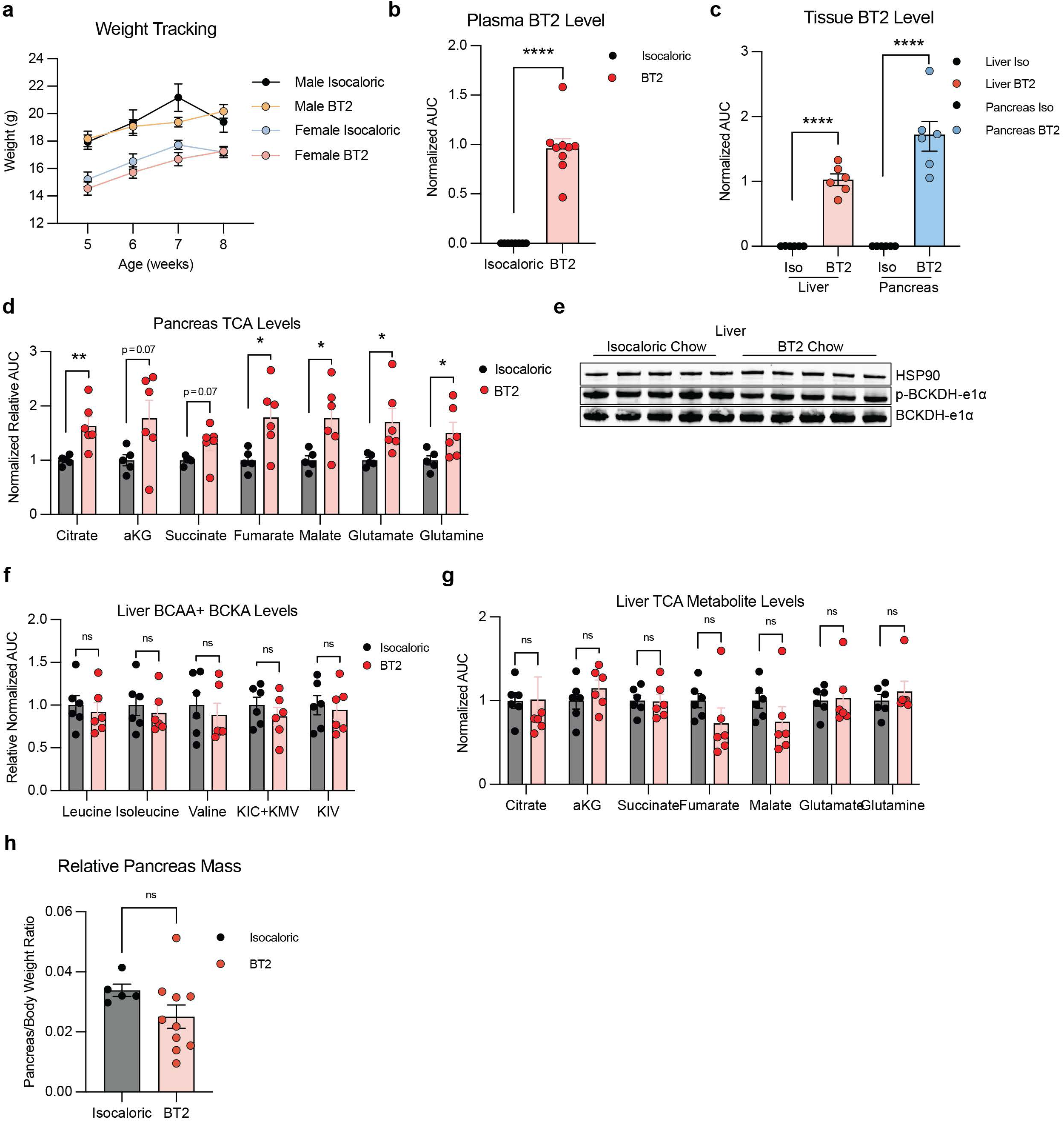
Further characterization of KC mice on BT2 chow. **a**, Animal weight recorded weekly of animals euthanized at 8 weeks old with cerulein administration at 6 ½ weeks and given BT2 chow or isocaloric control for 4 weeks starting at 4 weeks old. Data are shown separated by sex (n=6 male isocaloric chow, n=6 male BT2 chow, n=7 female isocaloric chow, n=6 BT2 chow). **b**, LC-MS was used to confirm the presence of BT2 in the plasma of animals after a 5 hour fast (n=8 isocaloric chow and n=9 BT2 chow). **c**, LC-MS was used to measure BT2 levels in the liver and pancreas of animals fed BT2 chow at fed state after 3 weeks on diet (n=5 Isocaloric chow and n=6 BT2 chow). **d**, LC-MS relative measurements of pancreatic TCA metabolites collected in a fed state (n=5 isocaloric chow and n=6 BT2 chow). **e**, Western for p-BCKDH-e1α and BCKDH-e1α protein measured in liver after 1 week on diet from animals mentioned in Fig. 6d (n=5 per genotype; HSP90 is the loading control). Animals were harvested in fed state. **f**, LC-MS relative measurements of liver BCAA and BCKA metabolites collected in a fed state (n=6 isocaloric chow and n=6 BT2 chow). **g**, LC-MS relative measurements of liver TCA metabolites collected in a fed state (n=6 isocaloric chowand n=6 BT2 chow). **h**, Relative pancreas mass measured by normalizing pancreas weight at endpoint to total body weight corresponding to Fig. 6f (n=5 Isocaloric chow and n=10 BT2 chow).Data are presented as mean± s.e.m. When two groups are compared, a two-tailed Student’s t-test was used, with significance defined as \**P* <0.05, \*\**P* <0.01 and \*\*\*\**P*<0.001. **b**, Plasma BT2 level *P*<0.0001. **c**, Liver BT2 level *P*<0.0001; Pancreas BT2 level *P*<0.0001. **d**, Citrate *P*=0.0079; aKG *P*=0.0676; Succinate *P*=0.0717; Fumarate *P*=0.0213; Malate *P*=0.0222; Glutamate *P*=0.0319; Glutamine *P*=0.0497.

## References

1. Mizrahi, J. D., Surana, R., Valle, J. W. & Shroff, R. T. Pancreatic cancer. The Lancet 395, 2008–2020 (2020).

2. American Cancer Society. Cancer Facts & Figures 2024. https://www.cancer.org/research/cancer-facts-statistics/all-cancer-facts-figures/2024-cancer-facts-figures.html.

3. Surveillance, Epidemiology, and End Results Program. Cancer of the Pancreas - Cancer Stat Facts. SEER https://seer.cancer.gov/statfacts/html/pancreas.html.

4. Stolzenberg-Solomon, R. Z., Schairer, C., Moore, S., Hollenbeck, A. & Silverman, D. T. Lifetime adiposity and risk of pancreatic cancer in the NIH-AARP Diet and Health Study cohort123. The American Journal of Clinical Nutrition 98, 1057–1065 (2013).

5. Rebours, V. et al. Obesity and Fatty Pancreatic Infiltration Are Risk Factors for Pancreatic Precancerous Lesions (PanIN). Clinical Cancer Research 21, 3522–3528 (2015).

6. Everhart, J. & Wright, D. Diabetes Mellitus as a Risk Factor for Pancreatic Cancer: A Meta-analysis. JAMA 273, 1605–1609 (1995).

7. Andersen, D. K. et al. Diabetes, Pancreatogenic Diabetes, and Pancreatic Cancer. Diabetes 66, 1103–1110 (2017).

8. Kirkegård, J. et al. Acute pancreatitis as an early marker of pancreatic cancer and cancer stage, treatment, and prognosis. Cancer Epidemiology 64, 101647 (2020).

9. Kim, H. S. et al. Incidence and risk of pancreatic cancer in patients with chronic pancreatitis: defining the optimal subgroup for surveillance. Sci Rep 13, 106 (2023).

10. Le Cosquer, G. et al. Pancreatic Cancer in Chronic Pancreatitis: Pathogenesis and Diagnostic Approach. Cancers 15, 761 (2023).

11. Zhang, A. M. Y. et al. Endogenous Hyperinsulinemia Contributes to Pancreatic Cancer Development. Cell Metabolism 30, 403–404 (2019).

12. Zhang, A. M. Y. et al. Effects of hyperinsulinemia on pancreatic cancer development and the immune microenvironment revealed through single-cell transcriptomics. Cancer & Metabolism 10, 5 (2022).

13. Zhang, A. M. Y. et al. Hyperinsulinemia acts via acinar insulin receptors to initiate pancreatic cancer by increasing digestive enzyme production and inflammation. Cell Metabolism 35, 2119–2135.e5 (2023).

14. Chung, K. M. et al. Endocrine-Exocrine Signaling Drives Obesity-Associated Pancreatic Ductal Adenocarcinoma. Cell 181, 832–847.e18 (2020).

15. Rossmeislová, L., Gojda, J. & Smolková, K. Pancreatic cancer: branched-chain amino acids as putative key metabolic regulators? Cancer Metastasis Rev 40, 1115–1139 (2021).

16. Eibl, G. et al. Diabetes Mellitus and Obesity as Risk Factors for Pancreatic Cancer. J Acad Nutr Diet 118, 555–567 (2018).

17. Blair, M. C., Neinast, M. D. & Arany, Z. Whole-body metabolic fate of branched-chain amino acids. Biochemical Journal 478, 765–776 (2021).

18. Katagiri, R. et al. Increased Levels of Branched-Chain Amino Acid Associated With Increased Risk of Pancreatic Cancer in a Prospective Case–Control Study of a Large Cohort. Gastroenterology 155, 1474–1482.e1 (2018).

19. Mayers, J. R. et al. Elevation of circulating branched-chain amino acids is an early event in human pancreatic adenocarcinoma development. Nat Med 20, 1193–1198 (2014).

20. Kamphorst, J. J. et al. Human pancreatic cancer tumors are nutrient poor and tumor cells actively scavenge extracellular protein. Cancer Res 75, 544–553 (2015).

21. Jiang, W. et al. Pancreatic stellate cells regulate branched-chain amino acid metabolism in pancreatic cancer. Ann Transl Med 9, 417 (2021).

22. Rossi, M. et al. Dietary intake of branched-chain amino acids and pancreatic cancer risk in a case–control study from Italy. British Journal of Nutrition 129, 1574–1580 (2023).

23. Li, J.-T. et al. BCAT2-mediated BCAA catabolism is critical for development of pancreatic ductal adenocarcinoma. Nat Cell Biol 22, 167–174 (2020).

24. Li, J.-T. et al. Diet high in branched-chain amino acid promotes PDAC development by USP1-mediated BCAT2 stabilization. National Science Review 9, nwab212 (2022).

25. Lei, M.-Z. et al. Acetylation promotes BCAT2 degradation to suppress BCAA catabolism and pancreatic cancer growth. Sig Transduct Target Ther 5, 1–9 (2020).

26. Neinast, M. D. et al. Quantitative Analysis of the Whole-Body Metabolic Fate of Branched-Chain Amino Acids. Cell Metabolism 29, 417–429.e4 (2019).

27. Carrer, A. et al. Acetyl-CoA Metabolism Supports Multistep Pancreatic Tumorigenesis. Cancer Discov 9, 416–435 (2019).

28. Hingorani, S. R. et al. Preinvasive and invasive ductal pancreatic cancer and its early detection in the mouse. Cancer Cell 4, 437–450 (2003).

29. Mayers, J. R. et al. Tissue of origin dictates branched-chain amino acid metabolism in mutant Kras-driven cancers. Science 353, 1161–1165 (2016).

30. Wang, X. & Proud, C. G. The mTOR Pathway in the Control of Protein Synthesis. Physiology 21, 362–369 (2006).

31. Westphalen, C. B. & Olive, K. P. Genetically Engineered Mouse Models of Pancreatic Cancer. Cancer J 18, 502–510 (2012).

32. Storz, P. Acinar cell plasticity and development of pancreatic ductal adenocarcinoma. Nat Rev Gastroenterol Hepatol 14, 296–304 (2017).

33. Kris, M. G., et al. Identification of driver mutations in tumor specimens from 1,000 patients with lung adenocarcinoma: The NCI’s Lung Cancer Mutation Consortium (LCMC). JCO 29, CRA7506–CRA7506 (2011).

34. Tang, Y. et al. Targeting KRASG12D mutation in non-small cell lung cancer: molecular mechanisms and therapeutic potential. Cancer Gene Ther 31, 961–969 (2024).

35. Yin, M. & Lei, Q.-Y. BCAT2–BCKDH metabolon maintains BCAA homeostasis. Nat Metab 4, 1618–1619 (2022).

36. Strauss, K. A., Puffenberger, E. G. & Carson, V. J. Maple Syrup Urine Disease. in GeneReviews® (eds. Adam, M. P. et al.) (University of Washington, Seattle, Seattle (WA), 1993).

37. Amaral, A. U. & Wajner, M. Pathophysiology of maple syrup urine disease: Focus on the neurotoxic role of the accumulated branched-chain amino acids and branched-chain α-keto acids. Neurochemistry International 157, 105360 (2022).

38. Liu, S. et al. Elevated branched-chain α-keto acids exacerbate macrophage oxidative stress and chronic inflammatory damage in type 2 diabetes mellitus. Free Radical Biology and Medicine 175, 141–154 (2021).

39. Sun, H. et al. Catabolic Defect of Branched-Chain Amino Acids Promotes Heart Failure. Circulation 133, 2038–2049 (2016).

40. Döppler, H. R., Liou, G.-Y. & Storz, P. Generation of Hydrogen Peroxide and Downstream Protein Kinase D1 Signaling Is a Common Feature of Inducers of Pancreatic Acinar-to-Ductal Metaplasia. Antioxidants 11, 137 (2022).

41. Döppler, H. R. & Storz, P. Macrophage-induced reactive oxygen species in the initiation of pancreatic cancer: a mini-review. Front. Immunol. 15, (2024).

42. Blair, M. C. et al. Branched-chain amino acid catabolism in muscle affects systemic BCAA levels but not insulin resistance. Nat Metab 5, 589–606 (2023).

43. Tso, S.-C. et al. Benzothiophene carboxylate derivatives as novel allosteric inhibitors of branched-chain α-ketoacid dehydrogenase kinase. J Biol Chem 289, 20583–20593 (2014).

44. East, M. P., Laitinen, T. & Asquith, C. R. M. BCKDK: an emerging kinase target for metabolic diseases and cancer. Nature Reviews Drug Discovery 20, 498–498 (2021).

45. Murashige, D. et al. Extra-cardiac BCAA catabolism lowers blood pressure and protects from heart failure. Cell Metabolism 34, 1749–1764.e7 (2022).

46. Yahsi, B. & Gunaydin, G. Immunometabolism – The Role of Branched-Chain Amino Acids. Front. Immunol. 13, (2022).

47. Lu, M. et al. Branched-chain amino acid catabolism promotes M2 macrophage polarization. Front. Immunol. 15, 1469163 (2024).

48. Yao, C., et al. Accumulation of branched-chain amino acids reprograms glucose metabolism in CD8+ T cells with enhanced effector function and anti-tumor response. Cell Reports 42, (2023).

49. Yang, Q., et al. BCKDK modification enhances the anticancer efficacy of CAR-T cells by reprogramming branched chain amino acid metabolism. Molecular Therapy 32, 3128– 3144 (2024).

50. Bowman, C. E. et al. Off-target depletion of plasma tryptophan by allosteric inhibitors of BCKDK. Mol Metab 97, 102165 (2025).

51. Acevedo, A. et al. The BCKDK inhibitor BT2 is a chemical uncoupler that lowers mitochondrial ROS production and de novo lipogenesis. J Biol Chem 300, 105702 (2024).

52. Roth Flach, R. J., et al. Small molecule branched-chain ketoacid dehydrogenase kinase (BDK) inhibitors with opposing effects on BDK protein levels. Nat Commun 14, 4812 (2023).

53. Filipski, K. J. et al. Discovery of First Branched-Chain Ketoacid Dehydrogenase Kinase (BDK) Inhibitor Clinical Candidate PF-07328948. J. Med. Chem. 68, 2466–2482 (2025).

54. White, P. J. et al. The BCKDH Kinase and Phosphatase Integrate BCAA and Lipid Metabolism via Regulation of ATP-Citrate Lyase. Cell Metabolism 27, 1281–1293.e7 (2018).

55. Heinemann-Yerushalmi, L. et al. BCKDK regulates the TCA cycle through PDC in the absence of PDK family during embryonic development. Dev Cell 56, 1182–1194.e6 (2021).

56. Cook, K. G., Bradford, A. P. & Yeaman, S. J. Resolution and reconstitution of bovine kidney branched-chain 2-oxo acid dehydrogenase complex. Biochem J 225, 731–735 (1985).

57. Lee, J. H. et al. Branched-chain amino acids sustain pancreatic cancer growth by regulating lipid metabolism. Exp Mol Med 51, 1–11 (2019).

58. Zhu, Z. et al. Tumour-reprogrammed stromal BCAT1 fuels branched-chain ketoacid dependency in stromal-rich PDAC tumours. Nat Metab 2, 775–792 (2020).

59. Demetriadou, C. et al. A nuclear branched-chain amino acid catabolism pathway controls histone propionylation in pancreatic cancer. 2025.04.23.650241 Preprint at 10.1101/2025.04.23.650241 (2025).

60. Xue, P. et al. BCKDK of BCAA Catabolism Cross-talking With the MAPK Pathway Promotes Tumorigenesis of Colorectal Cancer. EBioMedicine 20, 50–60 (2017).

61. Wang, Y. et al. BCKDK alters the metabolism of non-small cell lung cancer. Transl Lung Cancer Res 10, 4459–4476 (2021).

62. Tian, Q., et al. Phosphorylation of BCKDK of BCAA catabolism at Y246 by Src promotes metastasis of colorectal cancer. Oncogene 39, 3980–3996 (2020).

63. Xu, C. et al. BCKDK regulates breast cancer cell adhesion and tumor metastasis by inhibiting TRIM21 ubiquitinate talin1. Cell Death Dis 14, 1–11 (2023).

64. Ibrahim, S. L. et al. Inhibition of branched-chain alpha-keto acid dehydrogenase kinase augments the sensitivity of ovarian and breast cancer cells to paclitaxel. Br J Cancer 128, 896–906 (2023).

65. Abed, M. N. & Richardson, A. Inhibition of BCKDK increases the sensitivity of ovarian cancer cells to paclitaxel. European Journal of Cancer 69, (2016).

66. Biswas, D. et al. Inhibiting BCKDK in triple negative breast cancer suppresses protein translation, impairs mitochondrial function, and potentiates doxorubicin cytotoxicity. Cell Death Discov. 7, 1–12 (2021).

67. Bollinger, E. et al. BDK inhibition acts as a catabolic switch to mimic fasting and improve metabolism in mice. Molecular Metabolism 66, 101611 (2022).

68. DiMartino, S., Revelo, M. P., Mallipattu, S. K. & Piret, S. E. Activation of branched chain amino acid catabolism protects against nephrotoxic acute kidney injury. American Journal of Physiology-Renal Physiology 328, F152–F163 (2025).

69. He, Q.-Z., et al. 3,6-dichlorobenzo[b]thiophene-2-carboxylic acid alleviates ulcerative colitis by suppressing mammalian target of rapamycin complex 1 activation and regulating intestinal microbiota. World Journal of Gastroenterology 28, 6522–6536 (2022).

70. Zuo, X. et al. Multi-omic profiling of sarcopenia identifies disrupted branched-chain amino acid catabolism as a causal mechanism and therapeutic target. Nat Aging 5, 419–436 (2025).

71. Mullins, C. et al. Therapeutic Effects of BT2 in the Hippocampus of Alzheimer’s Disease Mouse Model. Curr Dev Nutr 6, 41 (2022).

72. Yeh, M.-C. et al. BT2 Suppresses Human Monocytic-Endothelial Cell Adhesion, Bone Erosion and Inflammation. JIR 14, 1019–1028 (2021).

73. Kawaguchi, Y. et al. The role of the transcriptional regulator Ptf1a in converting intestinal to pancreatic progenitors. Nat Genet 32, 128–134 (2002).

74. She, P. et al. Disruption of BCATm in mice leads to increased energy expenditure associated with the activation of a futile protein turnover cycle. Cell Metab 6, 181–194 (2007).

75. Jackson, E. L. et al. Analysis of lung tumor initiation and progression using conditional expression of oncogenic K-ras. Genes Dev. 15, 3243–3248 (2001).

76. Trefely, S., Ashwell, P. & Snyder, N. W. FluxFix: automatic isotopologue normalization for metabolic tracer analysis. BMC Bioinformatics 17, 485 (2016).

77. Agrawal, S. et al. El-MAVEN: A Fast, Robust, and User-Friendly Mass Spectrometry Data Processing Engine for Metabolomics. Methods Mol Biol 1978, 301–321 (2019).

78. Kantner, D. S. et al. Comparison of colorimetric, fluorometric, and liquid chromatography-mass spectrometry assays for acetyl-coenzyme A. Anal Biochem 685, 115405 (2024).

79. Snyder, N. W. et al. Production of stable isotope-labeled acyl-coenzyme A thioesters by yeast stable isotope labeling by essential nutrients in cell culture. Anal Biochem 474, 59–65 (2015).

80. Schindelin, J. et al. Fiji: an open-source platform for biological-image analysis. Nat Methods 9, 676–682 (2012).

81. Livak, K. J. & Schmittgen, T. D. Analysis of relative gene expression data using real-time quantitative PCR and the 2(-Delta Delta C(T)) Method. Methods 25, 402–408 (2001).

82. Chen, S., Zhou, Y., Chen, Y. & Gu, J. fastp: an ultra-fast all-in-one FASTQ preprocessor. Bioinformatics 34, i884–i890 (2018).

83. Patro, R., Duggal, G., Love, M. I., Irizarry, R. A. & Kingsford, C. Salmon provides fast and bias-aware quantification of transcript expression. Nat Methods 14, 417–419 (2017).

84. Gentleman, R. C. et al. Bioconductor: open software development for computational biology and bioinformatics. Genome Biology 5, R80 (2004).

85. Love, M. I. et al. Tximeta: Reference sequence checksums for provenance identification in RNA-seq. PLoS Comput Biol 16, e1007664 (2020).

86. Smedley, D. et al. BioMart – biological queries made easy. BMC Genomics 10, 22 (2009).

87. Blighe, K. PCAtools: Everything Princiapl Components Analysis. (2025).

88. Love, M. I., Huber, W. & Anders, S. Moderated estimation of fold change and dispersion for RNA-seq data with DESeq2. Genome Biology 15, 550 (2014).

89. Pantano, L., Hutchinson, J., Barrera, V. & Piper, M. DEGreport. (2025).

90. Hruban, R. H. et al. Pathology of Genetically Engineered Mouse Models of Pancreatic Exocrine Cancer: Consensus Report and Recommendations. Cancer Research 66, 95– 106 (2006).

91. DuPage, M., Dooley, A. L. & Jacks, T. Conditional mouse lung cancer models using adenoviral or lentiviral delivery of Cre recombinase. Nat Protoc 4, 1064–1072 (2009).

92. Crowe, A. R. & Yue, W. Semi-quantitative Determination of Protein Expression Using Immunohistochemistry Staining and Analysis: An Integrated Protocol. Bio-protocol 9, e3465 (2019).

